# Discovery of a First-in-Class SLIT2 Binder Disrupting the SLIT2/ROBO1 Axis via DNA-Encoded Library (DEL) Screening

**DOI:** 10.1101/2025.06.22.660907

**Authors:** Shaoren Yuan, Somaya A. Abdel-Rahman, Nelson García Vázquez, Hossam Nada, Laura Calvo-Barreiro, Katarzyna Kuncewicz, Moustafa T. Gabr

**Author notes:** Electronic Supplementary Information (ESI) available: Experimental procedures, and supplementary figures. For ESI or other electronic format see DOI: 10.1039/x0xx00000x.

## Abstract

The SLIT2/ROBO1 signaling axis plays a critical role in neural development, immune regulation, and tumor progression, including glioblastoma. However, small molecule inhibitors targeting this protein–protein interaction remain unexplored. Herein, we report the discovery and validation of **DEL-S1**, a first-in-class small molecule that binds to SLIT2 and disrupts its interaction with ROBO1. Using a DNA-encoded library (DEL) screen of 4.2 billion compounds, **DEL-S1** was identified and confirmed to bind SLIT2 via temperature-related intensity change (TRIC) assay. Functional inhibition of the SLIT2/ROBO1 complex by **DEL-S1** was demonstrated using a Time-Resolved Fluorescence Resonance Energy Transfer (TR-FRET) assay, yielding an IC_50_ of 68.8 ± 12.5 µM. Molecular docking and molecular dynamics (MD) simulations revealed key interaction hotspots at the SLIT2 binding interface and confirmed that **DEL-S1** impairs SLIT2/ROBO1 complex formation by inducing conformational rearrangements. **DEL-S1** exhibited favorable ADME properties, including satisfactory plasma and microsomal stability, low cytotoxicity, and minimal hERG liability. To facilitate structure–activity relationship (SAR) exploration, we designed and implemented a modular, one-pot synthetic route leveraging cyanuric chloride reactivity, enabling rapid derivatization of the triazine scaffold of **DEL-S1**. This strategy yielded structurally diverse analogs, including water-soluble carboxylate derivatives with preserved SLIT2/ROBO1 inhibitory activity. Together, this work establishes a novel chemical scaffold targeting SLIT2 and introduces a flexible synthetic platform to support further optimization toward therapeutic development.

## Introduction

Slit guidance ligands (SLITs) are glycoproteins that control axon guidance and neuronal migration, through binding to ROBO (Roundabout) receptors. ^1^In mammals, the SLIT family— comprising SLIT1, SLIT2, and SLIT3—binds to the immunoglobulin-like domain 1 (Ig1) of ROBO1 and ROBO2 via the D2 domain, a conserved leucine-rich repeat region.^2,3^ Beyond their classical role in axonal guidance, SLIT2/ROBO1 signaling is now recognized as a multifaceted regulator of physiological and pathological processes, including organ morphogenesis, cellular homeostasis, and tumor biology. This pathway not only orchestrates cellular motility and polarity but also modulates apoptosis, adhesion, and angiogenic responses, partly through downstream effectors like PI3K/Akt and various cytoskeletal adaptors.^3-5^

SLIT2/ROBO signaling mediates axonal repulsion in the developing central nervous system by activating intracellular pathways that remodel the cytoskeleton and redirect axonal extension.^6,7^ It also influences the migratory behavior of neurons^8^ and guides axonal growth and pathfinding.^9^ Beyond neurodevelopment, SLIT proteins exert diverse effects on immune cell trafficking. They function as chemoattractants for neutrophils while simultaneously repelling lymphocytes and dendritic cells, thereby creating cell type-specific patterns of immune cell positioning.^10-12^ In macrophages, SLIT2 modulates functional states by reducing macropinocytic activity and dampening the acquisition of pro-inflammatory or cytotoxic phenotypes.^13^ SLIT2 promotes vascular sprouting by directing tip cell alignment and migration, especially in retinal and skeletal tissues.^14-16^

In oncogenesis, SLIT2 functions in a context-dependent manner, exerting both tumor-promoting and tumor-restraining effects depending on the tissue and microenvironment. In several cancers—including colorectal, pancreatic, and osteosarcoma—SLIT2 is implicated in promoting invasive behaviors,^17^ supporting metastatic dissemination,^18-21^ and mediating resistance to therapy.^22^ However, in certain epithelial tumors such as breast and lung cancer, SLIT2/ROBO signaling is more often associated with growth inhibition and metastasis suppression.^23-25^ The role of this pathway in glioblastoma (GBM) remains particularly complex: while some studies indicate tumor-inhibitory properties,^26-28^ others highlight a pro-tumorigenic role,^29,30^ suggesting that its biological effects are highly dependent on the cellular context and tumor ecosystem. A growing body of evidence highlights SLIT2/ROBO signaling as a mechanism of immune suppression in the GBM tumor microenvironment (TME).^31^ Functional experiments have demonstrated that either silencing SLIT2 in glioma cells or blocking it systemically using a decoy receptor (ROBO1Fc) reduces TAM polarization and angiogenic gene signatures in preclinical GBM models.^31^

Despite these promising findings, clinical exploration of SLIT2/ROBO-targeted therapies remains limited. A protein therapeutic targeting this pathway, PF-06730512, was evaluated in patients with focal segmental glomerulosclerosis (FSGS), but the trial (NCT03448692) was terminated due to insufficient efficacy at tolerable doses.^32,33^ Furthermore, several ongoing trials (NCT03940820, NCT03941457, NCT03931720) are investigating ROBO1-directed CAR-NK cell therapies for solid tumors. Compared to biologics, small molecules offer several pharmacological advantages: reduced immunogenicity, improved control of exposure, and often favorable oral bioavailability.^34-37^ Their shorter systemic persistence also lowers the risk of extended on-target toxicities.^38,39^ Despite this, small molecule inhibitors of the SLIT2/ROBO1 axis are largely unexplored, representing a critical gap in the therapeutic landscape. The limitations of biologics—including restricted tissue penetration, manufacturing complexity, and potential immune activation— further underscore the need for small-molecule alternatives.^34-39^ Targeted small molecule-based therapies could provide a scalable and adaptable approach to disrupt SLIT2-mediated immune evasion and tumor progression, particularly in challenging cancers such as GBM.

We recently developed a high-throughput TR-FRET assay for the SLIT2/ROBO1 interaction, enabling for the first time the discovery of small molecule inhibitors targeting this pathway.^40^ Although the TR-FRET assay detects SLIT2/ROBO1 binding, directly targeting SLIT2 can disrupt upstream signaling, potentially yielding inhibitors with improved specificity and functional efficacy, thereby overcoming limitations of assays focused solely on protein-protein proximity. DNA-encoded library (DEL) technology has emerged as a transformative approach for the discovery of small molecule ligands capable of reversible target engagement, dramatically expanding the scope of chemical diversity beyond what is accessible through conventional high-throughput screening (HTS).^41-43^ By linking small molecules to unique DNA tags that serve as molecular barcodes, DEL platforms enable the construction and interrogation of libraries containing millions to billions of distinct entities.^41,42^ Screening is performed under pooled conditions, using minute amounts of the immobilized protein of interest. High-affinity binders are selectively retained on the target while nonspecific compounds are eliminated through washing. The retained molecules are then identified via DNA sequencing, which decodes the attached barcodes and enables rapid hit identification.^41-43^

In the present study, we applied DEL screening to discover novel small molecule binders targeting SLIT2, with the goal of disrupting its interaction with ROBO1. Our workflow for this study is presented in Figure 1. Starting from a DEL-derived hit, we confirmed its ability to bind SLIT2 using biophysical screening assays. We then demonstrated its capacity to inhibit the SLIT2/ROBO1 interaction using our recently reported TR-FRET assay^40^ for SLIT2/ROBO1 inhibition. Structural modeling of the SLIT2 binding interface and molecular dynamics (MD) simulations revealed key interaction hotspots, offering valuable guidance for future analog design and structure– activity relationship (SAR) studies. Remarkably, we evaluated the key pharmacokinetics (PK) and physicochemical properties of the DEL hit to inform future optimization efforts. Finally, we developed a synthetic route for the DEL hit, enabling the generation of an analog library to probe the structural features critical for SLIT2 binding. Given the limited number of successful applications of DEL technology to protein–protein interactions involved in immune regulation, our work underscores the potential of this platform to identify first-in-class SLIT2-targeted small molecules that inhibit SLIT2/ROBO1 interaction.

**Figure 1.**
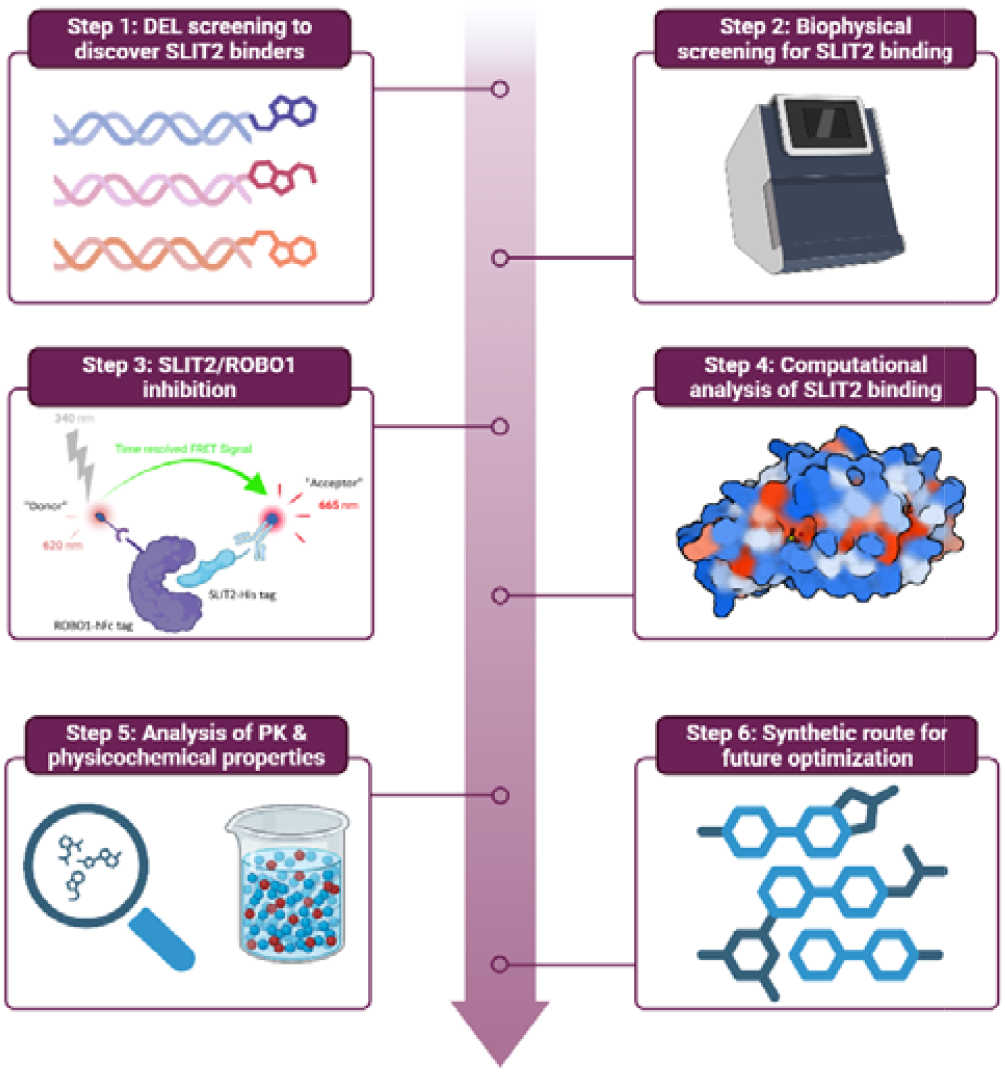
Workflow for the discovery of small molecule SLIT2 binders as inhibitoors of SLIT2/ROBO1 interaction.

## Results and discussion

### DEL screening

The DELopen kit is a publicly accessible DEL platform f**r**om WuXi AppTec that enables academic labs to perform affinity-based selections on diverse chemical matter withhout proprietary constraints. It includes structurally diverse libraries designed to broadly sample chemical space across multiple pharmacophores and scaffolds. A DEL screening of 4.2 billion compounds from the DELopen kit (WuXi AppTec) against the human SLIT2 protein resulted in the identification of 3,481 full-length compounds spanning 20 of the 27 included libraries (data not shown). Selection criteria included filtering out NTC signals (C4) from target conditions (C1, C2, and C3), applying an enrichment score threshold of 100, and prioriti**z**ing compounds that exhibited exclusive binding to the SLIT2 protein (A area) (Figure 2A). This multistep filtering pro**c**ess allowed us to eliminate false positives and emphasize selective SLIT2 engagement. Distinct enrichment patterns emerged from multiple libraries, and based on these trends (Figure 2B), along with the most prevalent chemotypes identified in the data analysis, the top small molecule SLIT2 binder (DEL-S1, Figure 2C) was selected for further investigation. Subsequently, we procured DEL-S1 from WuXi AppTec (purity >95%, see Figures S1-S5 for NMR, LCMS, mass spectra, HPLC purity, and chiral supercritical fluid chromatography (SFC)) to validate its potential to bind SLIT2 and inhibit key SLIT2 interactions.

**Figure 2.**
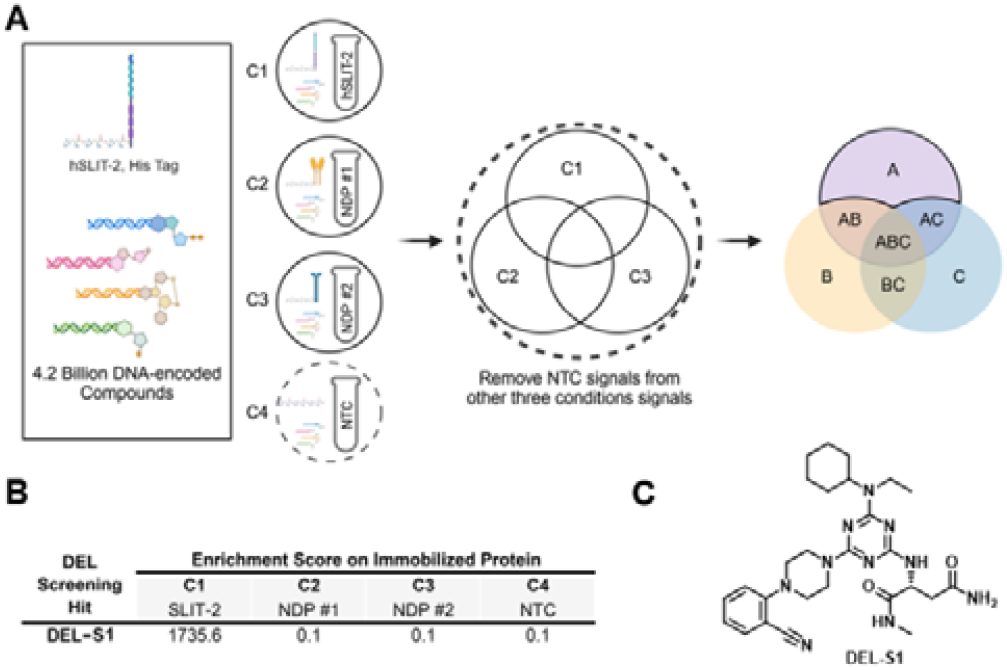
Identification of the top small molecule SLIT2 binder via DEL screening. **(A)** A total of 27 DNA-encoded libraries, encompassing 4.2 billion compounds, were screened against human SLIT2 protein immobilized on HisPur™ Ni-NTA Magnetic Beads (C1), along with two undisclosed human proteins immobilized on the same affinity matrix (C2, C3), and a non-target control (NTC, C4) consisting of the affinity matrix alone. Following affinity selection, bound small molecules were sequenced to decode their DNA tags. Compounds detected in area A, after filtering out NTC signals (C4) and applying selection criteria, were identified as potential SLIT2 binders. **(B)** Enrichment scores of **DEL-S1**, the top-selected small molecule binder for human SLIT2 following affinity selection. **(C)** Chemical structure of **DEL-S1**. Abbreviations: DEL, DNA-encoded library; NDP, non-disclosed protein.

### Biophysical validation

To validate **DEL-S1** as a first-in-class SLIT2-binding small molecule, we employed temperature-related intensity change (TRIC), a solution-phase biophysical technique that quantifies compound–protein interactions based on fluorescence changes under thermal gradients. Recombinant SLIT2 protein was fluorescently labeled and incubated with increasing concentrations of **DEL-S1**. The resulting thermophoretic signal demonstrated a clear, concentration-dependent response, consistent with a specific binding event (Figure 3). The observed TRIC signal suggests that **DEL-S1** engages SLIT2 in its native conformation, under near-physiological buffer conditions, without the need for immobilization or labeling of the compound. This is particularly valuable for protein–protein interaction targets like SLIT2, which often present shallow and dynamic binding surfaces that are challenging to interrogate using conventional biophysical approaches. Taken together, these results establish **DEL-S1** as a validated small molecule binder of SLIT2 and support its advancement into functional inhibition assays designed to probe its ability to disrupt the SLIT2/ROBO1 interaction.

**Figure 3.**
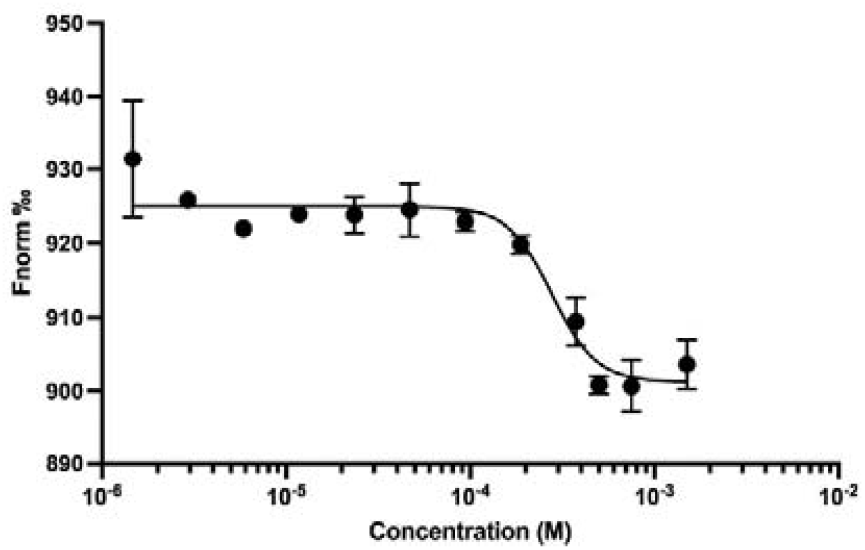
Dose-dependent curve for the binding of **DEL-S1** (12 assay points, 1.5 mMM – 1.5 µM, 2-fold dilution) to SLIT2 using TRIC.

### TR-FRET for SLIT2/ROBO1 inhibition

In order to validate **DEL-S1** as a SLIT2 binder capable of disrupting its functional interaction with ROBO1, we employed our previously optimized TR-FRET assay^40^ designed to quantify SLIT2/ROBO1 complex formation. In this system, d2-labeled SLIT2 and terbium-labeled ROBO1 were incubated in the presence of increasing concentrations of **DEL-S1**. In the absence of compound, a strong TR-FRET signal was observed, reflecting stable interaction between SLIT2 and ROBO1 (Figure 4). Titration of **DEL-S1** resulted in a concentration-dependdent decrease in TR-FRET efficiency, consistent with inhibitionn of the SLIT2/ROBO1 interaction. Nonlinear regression analysis of the dose–response curve yielded an IC_50_ value of 68.8 ± 12.5 μM (Figure 4), indicating the inhibitory activity of **DEL-S1** against this protein–protein interaction. Control experiments confirmed that the reduction in TR-FRET signal was not due to fluorescence quenching or compound interference with assay components. These findings establish **DEL-S1** as a functional small molecule inhibitor of the SLIT2/ROBO1 axis, underscoring its promise as a scaffold for therapeutic development in diseases such as glioblastoma.

**Figure 4.**
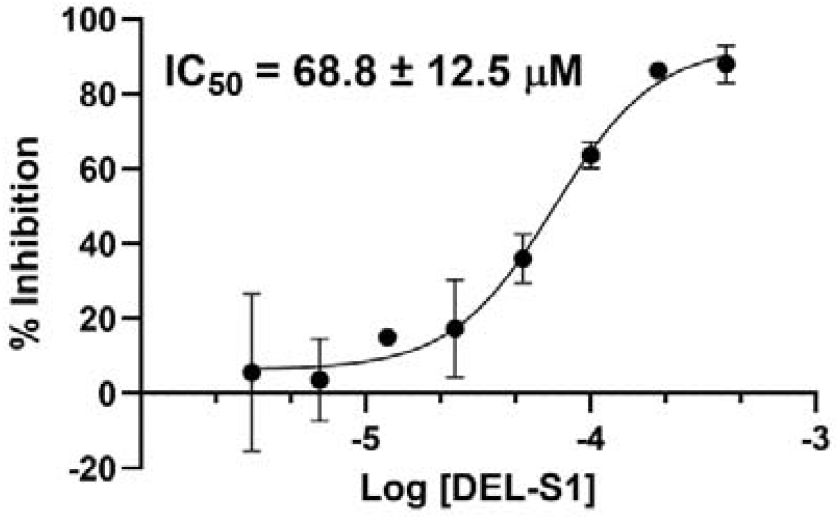
Dose-dependent curve for the inhibition of SLIT2/ROBO1 interaction by **DEL-S1** using TR-FRET assay. Error bars represent standard deviation (n=3).

### Computational analysis of the SLIT2 binding site

To better understand how our lead compound interacts with SLIT2 and guide future optimization efforts, we performed a detailed computational analysis of the SLIT2 binding site. This structural insight was aimed at identifying key molecular hotspots within the SLIT2 D2 domain that mediate its interaction with the ROBO1 Ig1 domain. The molecular interaction between ROBO1 and SLIT2 is mediated through a series of interactions between the Ig1 domain of ROBO1 and the concave face of the SLIT2 D2 domain. ^44^ This ROBO1/SLIT2 interaction involves two distinct binding regions (Figure 5A): an electrostatic interface formed by salt bridges (involving Ig1 residues Glu-72 and Lys-90 and SLIT2 D2 residues Arg-306 and Glu-304) and hydrogen bonds two hydrogen bonds (involving Asn-88 and Ser-75 of Ig1 and Arg-328 and Arg-287 of SLIT2 D2) between specific loops and strands, and a hydrophobic interface (involving Val-354, Ala-381, Leu-376, Leu-378, Leu-400, Ser-402, Tyr-404, His-426, and Ser-453)^44^ characterized by extensive apolar contacts between β-strand clusters (Figure 5B).

**Figure 5.**
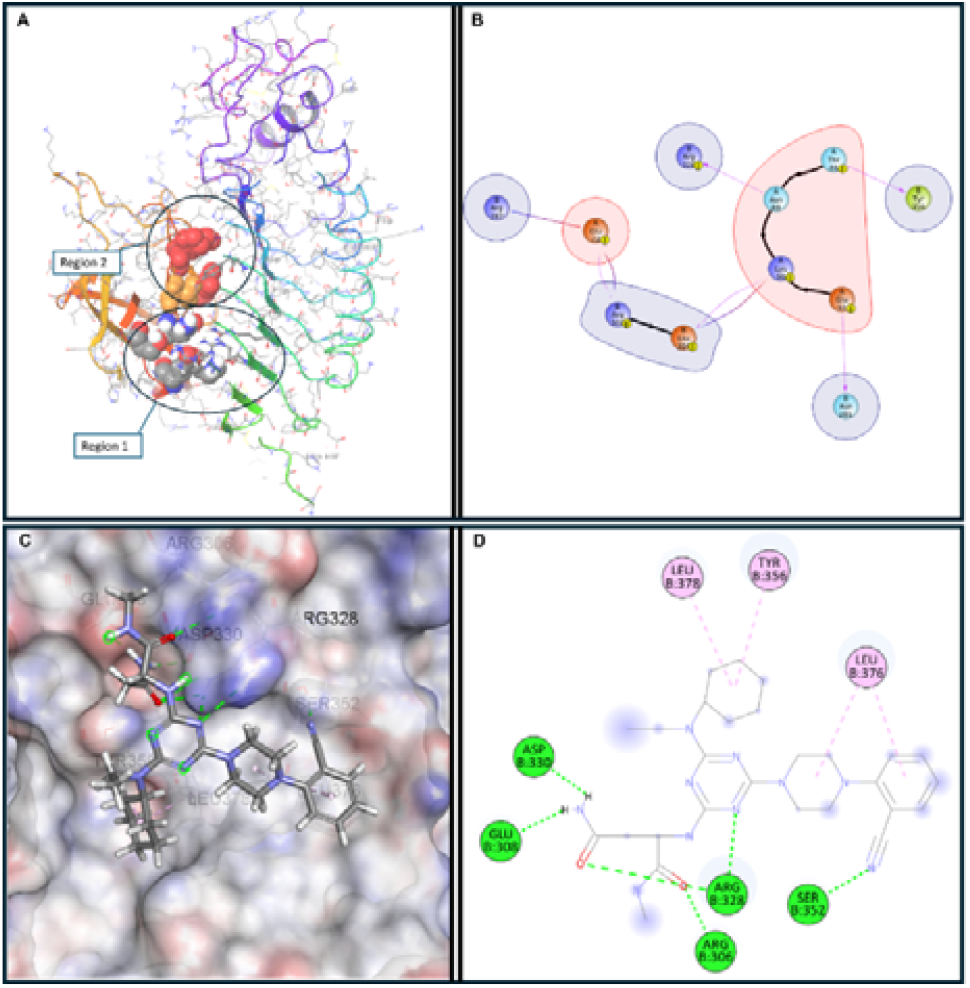
**(A)** The SLIT2/ROBO binding interface (PDB ID: 2V9T). **(B)** Interaction diagram illustrating key residue-level contacts within the native SLIT2/ROBO1 interface. **(C)** Three-dimensional binding pose of **DEL-S1** at the SLIT2 protein–protein interaction surface, showing occupancy of the ROBO1-binding pocket. **(D)** Two-dimensional interaction diagram of the **DEL-S1**/SLIT2 complex. Hydrogen bonds are depicted as dashed green lines, while π–π stacking interactions are shown in pink.

Induced Fit Docking (IFD) was employed to dock **DEL-S1** into the protein–protein interaction interface of SLIT2. The IFD protocol in Maestro (Schrödinger) was specifically used to account for the conformational flexibility of the SLIT2 binding site during ligand accommodation. The resulting binding pose of **DEL-S1** (Figure 5C-D) revealed that the compound established five hydrogen bonds with key residues in Region 1 of the SLIT2 interface, namely Arg306, Glu308, Arg328, Asp330, and Ser352. In addition to these polar interactions, SYT1 formed hydrophobic π–π stacking and van der Waals interactions with Tyr356, Leu376, and Leu378, which are part of Region 2 (Figure 5A). Collectively, these interactions suggest that **DEL-S1** is capable of engaging both critical regions of the SLIT2/ROBO binding interface, potentially disrupting their association and explaining the compound’s observed inhibitory activity.

To evaluate the effect of **DEL-S1** binding on the SLIT2/ROBO interaction, a ternary complex comprising SLIT2, **DEL-S1**, and ROBO was constructed and analyzed (Figure 6A-C). The ternary complex showed that **DEL-S1** binding to SLIT2 significantly disrupted the native SLIT2/ROBO1 protein–protein interface, leading to a notable reduction in both the complexity and extent of intermolecular contacts. The number of stabilizing contacts between SLIT2 and ROBO1 in the ternary complex was significantly reduced, with only a limited subset of residues remaining engaged (Figure 6C). In the **DEL-S1** -bound state, the interaction is primarily maintained through ROBO residues Glu-92, Glu-72, and Lys-90, interacting with SLIT2 residues Arg-287, Glu-304, and Asn-281. This represented a substantial loss of interface complexity compared to the native conformation, indicating that **DEL-S1** binding either induces conformational rearrangements in SLIT2 that misalign critical contact residues or directly competes with ROBO1 for overlapping binding sites.

**Figure 6.**
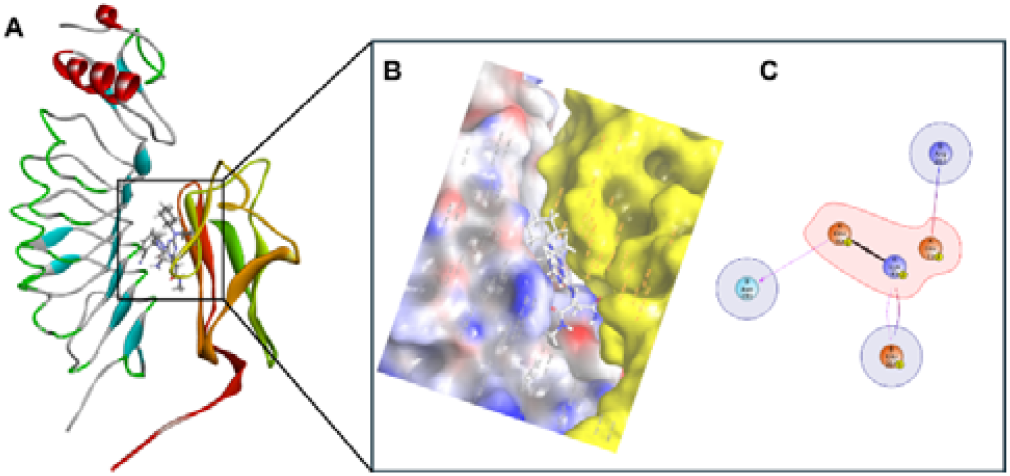
Effect of **DEL-S1** binding on the SLIT2–ROBO1 interaction. **(A)** Modeled ternary complex of **DEL-S1**-bound SLIT2–ROBO1, illustrating the spatial arrangement of all three components. **(B)** Three-dimensional surface representation of the ternary complex, highlighting the altered interaction interface upon **DEL-S1** binding. **(C)** 2D interaction diagram showing key residue-level contacts within the **DEL-S1**-bound SLIT2– ROBO1 interface, indicating the reduced and redistributed binding interactions compared to the native complex.

A 100 ns MD simulation was performed to assess the stability of the predicted **DEL-S1**/SLIT2 complex and its effect on the SLIT2/ROBO interaction. Throughout the MD simulation, **DEL-S1** remained stably bound to SLIT2 and was able to consistently maintain the hydrogen bond interaction with Arg328. In contrast, the distance between SLIT2 and ROBO1 progressively increased (Figure 7) over the course of the simulation which resulted in hindering ROBO1 from forming its native and residue-specific contacts within the canonical SLIT2/ROBO1 interface. Together, these observations explain the observed inhibitory activity of **DEL-S1** on the SLIT2/ROBO1 interaction.

**Figure 7.**
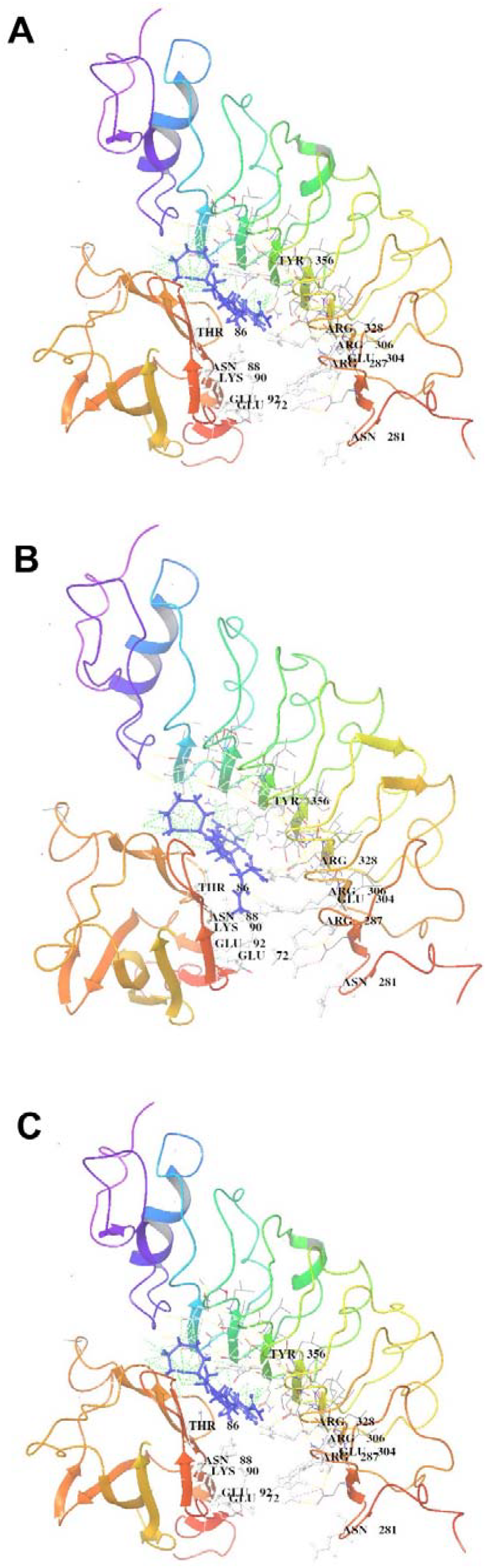
Conformational changes in the SLIT2–ROBO1 ternary complex during a 100 ns molecular dynamics simulation. **(A)** Snapshot of the initial structure of the ternary complex. **(B)** Intermediate structure at 50 ns showing partial separation between SLIT2 and ROBO1. **(C)** Final structure at 100 ns, highlighting a further increase in the distance between SLIT2 and ROBO1, suggesting progressive weakening of their interaction.

### Assessment of the key PK and physicochemical properties of DEL-S1

The development of small molecule therapeutics for cancer, particularly brain cancers, requires not only inhibition of disease-relevant pathways but also favorable PK and safety profiles that support effective drug delivery to the central nervous system (CNS). To evaluate the suitability of **DEL-S1** as a preliminary hit for CNS-targeted therapy, we performed a comprehensive in vitro ADME (absorption, distribution, metabolism, and excretion) assessment using standard zed assay protocols (Table 1).

**Table 1.** In vitro PK profile of DEL-S1.

| PK property | Value |
| --- | --- |
| LogD7.4<sup>a</sup> | 1.63 |
| Kinetic Solubility 1% DMSO/PBS [$\mu\text{M}$]<sup>b</sup> | 246 |
| PAMPA permeability [cm/s] | $2.1 \times 10^{-6}$ |
| PPB human [%]<sup>c</sup> | $88.2 \pm 3.16$ |
| $t_{1/2}$ [h] mouse liver microsomes<sup>c</sup> / $\text{CL}_{\text{int}}$<sup>d</sup> | $1.9 \pm 0.4$ / $18 \pm 1.4$ |
| $t_{1/2}$ [h] human liver microsomes<sup>c</sup> / $\text{CL}_{\text{int}}$<sup>d</sup> | $2.5 \pm 0.3$ / $22 \pm 1.2$ |
| $t_{1/2}$ [h] mouse plasma | $1.7 \pm 0.2$ |
| $t_{1/2}$ [h] human plasma | $2.1 \pm 0.3$ |
| $t_{1/2}$ [h] PBS | >4 |
| $\text{IC}_{50}$ against hERG [$\mu\text{M}$] | >100 |

<sup>a</sup>LogD (pH = 7.4) the distribution coefficient at physiological pH 7.4. <sup>b</sup>Kinetic solubility measured in 1% DMSO/PBS (pH 7.4) by UV-vis spectrophotometry (n=3). <sup>c</sup>Values are mean ± SD (n=3). <sup>d</sup>Intrinsic clearance expressed in $\mu\text{L}/(\text{min} \times \text{mg})$ protein. Values are mean ± SD (n=3).

Kinetic solubility of **DEL-S1** was measured at 246 µM, demonstrating acceptable aqueous solubility under physiologically relevant conditions. This parameter is particularly important for CNS-active drugs, as it enables sufficient free drug concentration in systemic circulation. This was further supported by its balanced logP of 1.63 (Table 1). The compound’s PAMPA (Parallel Artificial Membrane Permeability Assay) permeability was 2.1⍰× ⍰ 10^−6^ cm/s, consistent with moderate passive diffusion across lipid membranes, likely due to its high polarity. Plasma protein binding (PPB) of **DEL-S1** was measured at 88.2%, indicating a substantial proportion of the compound is bound in circulation, while still allowing a pharmacologically active free fraction.

**DEL-S1** exhibited moderate stability in both mouse and human plasma, with half-lives (t_1_/_2_) of 1.7⍰± ⍰ 0.2 h and 2.1⍰± ⍰ 0.3 h, respectively (Table 1). Microsomal stability assays revealed half-lives of 1.9⍰± ⍰ 0.4 h in mouse and 2.5⍰± ⍰ 0.3 h in human liver microsomes, indicating that **DEL-S1** is not rapidly metabolized and may maintain systemic exposure following administration. The intrinsic clearance (Cl_int_) was determined to be 1811±rn1.4 mL/min/mg (mouse) and 22⍰± ⍰ 1.2 mL/min/mg (human), consistent with moderate hepatic metabolism and favorable bioavailability characteristics.

To evaluate off-target cytotoxicity, **DEL-S1** was tested at 50 µM for 72 hours in three representative non-cancerous human cell lines: normal human astrocytes (NHA), human brain microvascular endothelial cells (hBMECs), and HepG2 hepatocytes. As shown in Figure S6, **DEL-S1** had minimal cytotoxic effects across all three cell types. Cell viability remained above 95% in NHA, hBMECs, and HepG2 cells, supporting a favorable safety profile. In vitro hERG liability assessment showed an IC_50_ > 100 µM, suggesting a low risk of cardiotoxicity at therapeutically relevant concentrations (Table 1). Collectively, these findings demonstrate that **DEL-S1** exhibits physicochemical and ADME properties compatible with small molecule drug development. Its balanced metabolic stability, aqueous solubility, passive permeability, and low off-target cytotoxicity render it a promising lead candidate for further hit-to-lead optimization studies.

### Synthetic route for optimization studies

To generate a focused analog library of **DEL-S1** for future SAR studies, we employed a modular synthetic strategy (Scheme 1) leveraging the electrophilic reactivity of cyanuric chloride (**1**), a highly reactive molecule featuring three chlorine atoms that serve as replaceable sites. These chlorine atoms can be sequentially substituted by various nucleophiles under controlled conditions with sequential steps, making cyanuric chloride a versatile scaffold for creating a wide range of triazine derivatives. By adjusting reaction conditions, such as temperature and the order of nucleophile addition, sequential nucleophilic substitutions are achieved in one-pot.

As shown in Scheme 1, amino acid esters were selected as the initial nucleophiles in the reaction sequence, driven by their relatively low nucleophilicity and limited solubility, necessitating the use of the most reactive electrophilic site on the cyanuric chloride scaffold. Conversely, piperazine derivatives, characterized by their high nucleophilic strength, were designated as the final nucleophiles to engage with the mono-chlorinated intermediates, facilitating complete substitution. This sequential arrangement can be adapted based on the nucleophilic properties of the reagents; for instance, the incorporation of highly nucleophilic secondary or primary amines could be adapted as the third reagent added in the synthesis. This step-by-step substitution enables the modular attachment of different substituents to the triazine core, producing a library of compounds (**2**, Scheme 1) tailored for SAR studies.

**Scheme 1.**
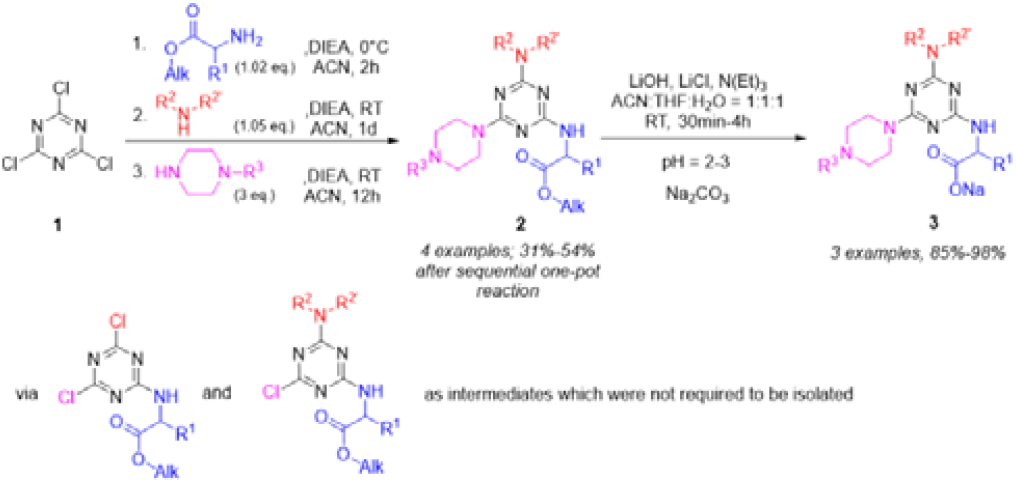
Synthesis of **DEL-S1** derivatives **(2)** via sequential one-pot nucleophilic aromatic substitution and selective hydrolysis products **(3)**.

This synthetic platform supports flexible adaptation based on the nucleophilic profile of the input reagents, allowing for rapid diversification of the triazine scaffold with both hydrophobic and polar substituents. As shown in Figure 8, the synthesized compounds (**2a-d**) included changes in three key fragments of the initial hit (**DEL-S1**). To improve aqueeous solubility and PK properties, selected analogs underwent ester hydrolysis (Scheme 1) to yield corresponding carboxylic aacid derivatives (**3**). This modification remarkably enhanced the solubility of the resulting **3a, 3b**, and **3d** compounds (Figuree 8), addressing a key limitation in early-stage drug candidates.

**Figure 8.**
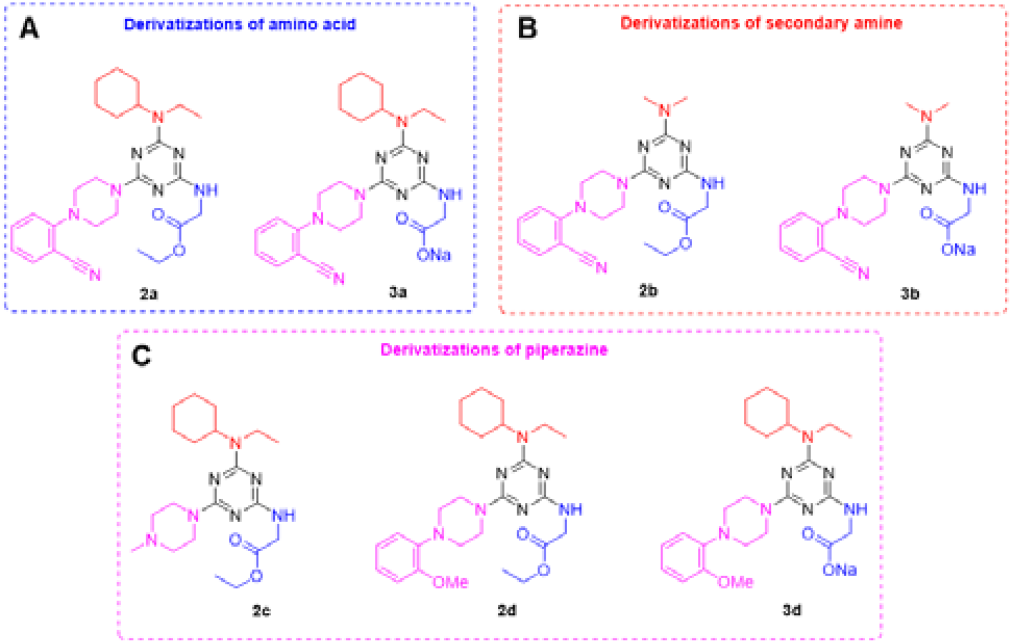
Chemical structures of the synthesized **DEL-S1** derivatives. **(A)** the derivatizations of amino acid; **(B)** the derivatizations of secondary amine; and **(C)** the derivatizations of piperazine.

Preliminary evaluation of the synthesized compounds indicated that the ester derivatives (**2a-d**) exhibited limited solubility under the SLIT2/ROBO1 TR-FRET assay conditioons, precluding their functional assessment. In contrast, corresponding sodium carboxylates (**3a, 3b**, and the **3d**) demonstrated improved solubility, allowing for poteency evaluation. Notably, compounds **3a** and **3d** possesssed SLIT2/ROBO1 inhibitory activity, with IC_50_ values of 225.4 µM and 238.8 µM, respectively. However, compound **3b** laccked inhibitory SLIT2/ROBO1 activity. Although none of the synthesized derivatives exceeded the potency of **DEL-S1**, the preservation of inhibitory activity in select analogs confirms the tractability of the **DEL-S1** chemotype and supportss its potential for further structural optimization. This streamlined synthetic strategy enabled the efficient generation oof a structurally diverse library, laying a solid foundation for futture SAR studies aimed at enhancing both potency and pharmacokinetic properties.

## Conclusions

In summary, we report the discovery and validation of **DEL-S1** as the first small-molecule binder of SLIT2 capable of disruptingg the SLIT2/ROBO1 interaction—a protein–protein interface long considered undruggable. Using a DEL screen of over 4 billion compounds, **DEL-S1** emerged as a selective SLIT2 ligand and was biophysically and functionally validated using TRIC and TR-FFRET assays. Computational modeling and molecular dynammics simulations revealed its binding mode and mechanistic disruptioon of the SLIT2/ROBO1 complex. To accelerate SAR exploration, we established a modular, one-pot synthetic route leveraging cyanuric chloride chemistry, enabling rapid generation of DEL-S1 analogs. This platform yielded water-soluble derivatives with SLIT2/ROBO1 inhibitory activity, demonstrating the tractability of the chemotype. Moreover, DEL-S1 showed favorable ADME properties, low cytotoxicity, and low hERG liability, underscoring its potential as a lead for CNS-targeted therapies. Collectively, this work not only identifies a novel chemical probe for the SLIT2/ROBO1 axis but also provides a versatile synthetic toolkit and roadmap for developing next-generation immunomodulatory therapeutics in oncology.

### Experimental

#### DEL Screening

A DEL comprising 4.2 billion compounds, supplied by WuXi AppTec, was utilized to screen for interactions with the human slit homolog 2 (SLIT2) protein. Prior to conducting affinity selection, a protein capture assay was performed to confirm the stable immobilization of SLIT2 protein (Cat. # SL2-H52H6, ACROBiosystems, Newark, DE, USA) onto the affinity matrix. This preliminary validation led to the optimization of the selection buffer, which consisted of 1x PBS, 0.05% Tween-20, 0.1 mg/mL sssDNA, and 10 mM imidazole. The elution buffer was formulated with 1x PBS and 0.05% Tween-20. Each round of selection employed 5 µg of SLIT2 protein in conjunction with HisPur™ Ni-NTA Magnetic Beads (Cat. # 88831, Thermo Fisher, Waltham, MA, USA) as the affinity matrix.

Affinity selection, encompassing both protein immobilization and subsequent selection rounds, was executed in accordance with the manufacturer’s protocol. The DELopen kit included four identical sets of 27 distinct libraries, collectively covering 4.2 billion compounds. One set was allocated for SLIT-2 screening, while two others were used for screening different human proteins (undisclosed). The fourth set functioned as a non-target control (NTC), using HisPur™ Ni-NTA Magnetic Beads without protein to assess nonspecific binding.

Following the second selection round, the heated elution sample underwent a self-QC step, ensuring the concentration of selected DEL molecules remained below 5×10^8^ copies (standard control) and free from cross-contamination. This post-selection sample was submitted to WuXi AppTec, where it passed sequencing quality control before advancing to next-generation sequencing (NGS) for decoding the DNA-linked small molecules. Data analysis revealed the top-binding compound. WuXi AppTec disclosed the structure to the authors, and upon request, synthesized a tag-free version for subsequent investigation.

#### TRIC assay

Dose response was performed on a Dianthus (NanoTemper, Munich, Germany) instrument, using the Temperature Related Intensity Change (TRIC) technology. 50 nM of hSLIT2-His labeled with RED-tris-NTA 2nd generation dye (NanoTemper, Munich, Germany) in assay buffer (10mM HEPES, 150mM NaCl, pH 7.4, 0.05% Tween20, 2% DMSO) was incubated with ligand (12 assay points, 1.5 mM – 1.5 µM, 2-fold dilution) for 20 minutes. TRIC measurements were conducted at ∼68% laser excitation, determined by instrument. The binding curve was obtained using GraphPad Prism 10.

#### TR-FRET assay

The SLIT2/ROBO1 TR-FRET assay was performed as we previously reported.^40^

#### Computational study

The crystal structure of the complex between the second leucine-rich repeat (LRR) domain of SLIT2 and the first immunoglobulin (Ig1) domain of ROBO1 (PDB ID: 2V9T) was retrieved from the Protein Data Bank. Structural preparation was carried out using the Protein Preparation Wizard module in Maestro (Schrödinger Suite, v2021.2) under default settings. This process involved the removal of water molecules and crystallographic artifacts, addition of any missing side chains and hydrogen atoms, and assignment of protonation states consistent with physiological pH (7.4). The prepared protein complex was then subjected to restrained energy minimization using the conjugate gradient algorithm with the OPLS-2005 force field to achieve a stable, low-energy conformation suitable for docking studies. SYT1 was prepared using the LigPrep module in Maestro with default parameters. This included generation of low-energy conformers, assignment of appropriate protonation states at pH 7.4, and energy minimization to ensure accurate geometry. Molecular docking of SYT1 into the SLIT2 binding pocket was performed using the Induced Fit Docking (IFD) workflow in Maestro which allows for receptor flexibility during ligand accommodation. Default parameters were used throughout the IFD protocol to predict the most favorable binding pose within the protein–protein interface. Discovery studio visualizer was employed to demonstrate the 2D and 3D interaction diagrams of SYT1 in complex with SLIT2. MD simulation was carried out and analyzed using DESMOND software under default conditions using the same methodology as our previous work.^45^

#### In vitro PK studies

The These experiments were conducted following our previously reported methods.^45^ The evaluations included LogD7.4 determination, microsomal stability, kinetic solubility, and cytotoxicity profiling across multiple cell lines. Solubility was assessed using UV–visible spectrophotometry, while cell viability was measured using the PrestoBlue assay.

#### Synthetic chemistry

All reactions were performed under protection of N_2_ in oven-dried glassware unless water was applied as solvent. Chromatographic purification was performed as flash chromatography with Combi-Flash^®^ Rf+ UV-VIS MS COMP using RediSep^®^ Silver normal phase silica gel columns and solvents indicated as eluent with default pressure. Fractions were collected based on UV absorption at 254 nm and/or 280 nm. Analytical thin layer chromatography (TLC) was performed on Whatman ^®^ TLC silica gel UV254 (250 µm) TLC aluminum plates. Visualization was usually accomplished with UV light (254 nm and 365 nm). Proton and carbon nuclear magnetic resonance spectra (H NMR and C NMR) were recorded on a Bruker 500 MHz spectrometer with solvent resonances as the internal standard (^1^H NMR: CDCl_3_ at 7.26 ppm, DMSO-d6 at 2.50 ppm, Acetone-d6 at 2.05 ppm; ^13^C NMR: CDCl_3_ at 77.0 ppm, DMSO-d_6_ at 39.52 ppm, Acetone-d6 at 29.84 ppm and 206.26 ppm). ^1^H NMR data are reported as follows: chemical shift (ppm), multiplicity (s = singlet, d = doublet, dd = doublet of doublets, dt = doublet of triplets, ddd = doublet of doublet of doublets, t = triplet, m = multiplet, br = broad), coupling constants (Hz), and integration. The NMR spectra of some sodium carboxylate final products might contain small amount of triethylamine hydrochloride during the work-up procedure. Occasionally, the ^13^C signals of triazine carbons might not be visible. Mass spectra were obtained through LC-MS on Waters Acquity UPLC^®^ and Zspray™.

#### General Procedure A: the sequential one-pot synthesis of 1,3,5-triazine derivatives 2

A 50 mL oven-dried round-bottom flask equipped with a stir bar was purged with dry N_2_. Cyanuric chloride (400 mg, 2.17 mmol, 1 eq.) was added to the flask, followed by anhydrous acetonitrile (5 mL). The mixture was cooled to 0°C. In a separate vial, amino acid alkyl ester (2.21 mmol, 1.02 eq.) and N,N-diisopropylethylamine (DIEA, 323 mg, 435 μL, 2.5 mmol, 1.15 eq.) were dissolved or suspended in anhydrous acetonitrile (5 mL). (*Note: The amount of DIEA was doubled when amino acid alkyl ester HCl salt was used*.) This mixture was added dropwise to the flask over 10 minutes at 0°C. After complete addition, the reaction was maintained at 0°C for an additional 2 hours.

In a second separate vial, secondary amine (2.28 mmol, 1.05 eq.) and DIEA (323 mg, 435 μL, 2.5 mmol, 1.15 eq.) were dissolved in anhydrous acetonitrile (2 mL). The resulting solution was added rapidly to the flask. The reaction mixture was then warmed to room temperature and stirred for 24 hours. Subsequently, a mixture of piperazine derivatives (6.51 mmol, 3 eq.) and DIEA (323 mg, 435 μL, 2.5 mmol, 1.15 eq.) in anhydrous acetonitrile (2 mL) was added rapidly to the flask. The reaction was stirred at room temperature for 12 additional hours, with complete consumption of the monochloride intermediate monitored by TLC.

Upon reaction completion, the mixture was poured into water, and the aqueous layer was acidified to pH 3–4 to neutralize unreacted piperazine. The aqueous layer was then extracted with ethyl acetate (2 × 100 mL). The combined organic layers were washed with brine, dried over Na_2_SO_4_, filtered, and concentrated under reduced pressure to obtain crude product, which was absorbed onto a plug of silica gel and purified by chromatography.

#### General Procedure B: the synthesis of 3 via the selectively hydrolysis of 2

To a 20 mL scintillation vial charged a stir bar, **2** (0.2 mmol, 1 eq.), was added LiOH (14 mg, 0.6 mmol, 3 eq.) and LiCl (85 mg, 2 mmol, 10 eq.). A mixture of water (1.3 mL), acetonitrile (1.3 mL) and THF (1.3 mL) was added to the vial, followed by N(Et)_3_ (41 mg, 56mL, 0.4 mmol, 2 eq.). The reaction mixture was stirred at room temperature for 30 mins, poured into water, and the aqueous layer was acidified to pH 2–3. The aqueous layer was then extracted with ethyl acetate or 30% IPA in DCM (2 × 50 mL). The combined organic layers were washed with brine, dried over Na_2_SO_4_, filtered, and solvent was removed under reduced pressure to obtain the carboxylic acid, which can further react with 0.1M Na_2_CO_3_ to produce the sodium carboxylate product 3.

Compound **2a** (ethyl (4-(4-(2-cyanophenyl)piperazin-1-yl)-6-(cyclohexyl(ethyl)amino)-1,3,5-triazin-2-yl)glycinate). **2a** was obtained (541 mg, 1.10 mmol, 51%) as light-yellow powder by following general procedure A with glycine ethyl ester hydrochloride (309 mg), N-ethylcyclohexanamine (290 mg) and 2-(piperazin-1-yl)benzonitrile (1.22 g), and the purification was accomplished by chromatography with 10%-50% EtOAc in hexanes. ^1^**H NMR** (500 MHz, CDCl_3_) δ 7.58 (dd, J = 7.7, 1.8 Hz, 1H), 7.48 (ddd, *J* = 9.2, 7.5, 1.7 Hz, 1H), 7.04 – 6.98 (m, 2H), 5.23 (s, 1H, br), 4.45 (*t, J* = 12.4 Hz, 1H), 4.20 (q, *J* = 7.2 Hz, 2H), 4.10 (d, *J* = 5.8 Hz, 2H), 3.99 – 3.93 (m, 4H), 3.43 (q, *J* = 6.9 Hz, 2H), 3.22 – 3.17 (m, 4H), 1.87 – 1.62 (m, 5H), 1.49 – 1.31 (m, 4H), 1.28 (*t, J* = 7.1 Hz, 3H), 1.18 – 1.07 (m, 4H). ^13^**C NMR** (126 MHz, CDCl_3_) δ 170.95, 165.72, 164.85, 164.42, 155.74, 134.33, 133.77, 121.89, 118.74, 118.35, 106.19, 60.96, 54.47, 51.55, 43.18, 43.04, 37.15, 30.95, 26.13, 25.76, 15.21, 14.20. LC-MS (ES+): m/z 493.95 [M+H]^+^

Compound **3a** (sodium (4-(4-(2-cyanophenyl)piperazin-1-yl)-6-(cyclohexyl(ethyl)amino)-1,3,5-triazin-2-yl)glycinate). **3a** was obtained (91 mg, 0.187 mmol, 92%) as yellow powder by following general procedure B with **2a** (100 mg). ^1^**H NMR** (500 MHz, Acetone) δ 7.57 (dd, J = 7.6, 1.5 Hz, 1H), 7.54 – 7.47 (m, 1H), 7.04 (t, J = 7.6 Hz, 1H), 7.01 – 6.96 (m, 1H), 6.26 (s, 1H, br), 4.04 (d, J = 7.2 Hz, 2H), 3.86 (s, 4H), 3.43 – 3.32 (m, 2H), 3.07 (s, 4H), 1.77 – 1.53 (m, 5H), 1.47 – 1.24 (m, 5H), 1.09 (t, J = 7.1 Hz, 4H). ^13^**C NMR** (126 MHz, Acetone) δ 177.89, 166.59, 166.24, 165.66, 156.77, 135.10, 135.01, 122.92, 120.06, 119.04, 107.00, 52.49, 46.35, 44.12, 37.81, 31.94, 27.14, 26.67, 16.35, 16.10. LC-MS (ES+): m/z 465.44 [M-Na+2H]^+^

Compound **2b** (ethyl (4-(4-(2-cyanophenyl)piperazin-1-yl)-6-(dimethylamino)-1,3,5-triazin-2-yl)glycinate). **2b** was obtained (305 mg, 0.743 mmol, 34%) as light pink solid by following the first step of general procedure A with glycine ethyl ester hydrochloride (309 mg). Instead of adding dimethylamine motif as the second step, 2-(piperazin-1-yl)benzonitrile (426 mg, 2.28 mmol, 1.05 eq.) was added with DIEA. After 12 hours, the acetonitrile solution of dimethylamine hydrochloride (1.04 g, 10.85 mmol, 5 eq.) with DIEA (1.4 g, 10.85 mmol, 5 eq.) was added to the reaction, which was stirred at room temperature for 2 additional hours, with complete consumption of the monochloride intermediate monitored by TLC. Purification was accomplished by chromatography with 10%-50% ethyl acetate in hexanes. ^1^**H NMR** (500 MHz, CDCl_3_) δ 7.59 (dd, *J* = 7.7, 1.8 Hz, 1H), 7.50 (ddd, *J* = 8.9, 7.6, 1.8 Hz, 1H), 7.04 (*t, J* = 7.6 Hz, 1H), 7.00 (d, *J* = 8.2 Hz, 1H), 4.22 (q, *J* = 7.1 Hz, 2H), 4.13 (d, *J* = 5.8 Hz, 2H), 4.03 – 3.97 (m, 4H), 3.24 – 3.18 (m, 4H), 3.12 (s, 6H, br), 1.28 (*t, J* = 7.2 Hz, 3H). ^13^**C NMR** (126 MHz, CDCl_3_) δ 168.31, 155.56, 134.37, 133.84, 122.17, 118.83, 118.27, 106.41, 61.14, 51.51, 43.45, 43.00, 36.19, 14.24. LC-MS (ES+): m/z 411.88 [M+H]^+^

Compound **3b** (sodium (4-(4-(2-cyanophenyl)piperazin-1-yl)-6-(dimethylamino)-1,3,5-triazin-2-yl)glycinate). **3b** was obtained (79 mg, 0.195 mmol, 98%) as yellow powder by following general procedure B with **2b** (82 mg). ^1^**H NMR** (500 MHz, CDCl_3_) δ 7.48 (d, *J* = 7.8 Hz, 1H), 7.39 (*t, J* = 8.0 Hz, 1H), 6.95 (q, *J* = 7.5 Hz, 1H), 6.86 (d, *J* = 8.5 Hz, 1H), 3.99 (s, 2H, br), 3.85 (s, 4H, br), 3.14 – 3.02 (m, 4H), 2.97 (s, 6H, br).^13^ **C NMR** (126 MHz, CDCl_3_) δ 176.30, 155.50, 134.19, 133.86, 121.85, 118.85, 118.34, 105.91, 57.98, 51.59, 43.34, 36.47. LC-MS (ES+): m/z 383.82 [M-Na+2H]^+^

Compound **2c** (ethyl (4-(cyclohexyl(ethyl)amino)-6-(4-methylpiperazin-1-yl)-1,3,5-triazin-2-yl)glycinate). **2c** was obtained (423 mg, 1.04 mmol, 48%) as light yellow amorphous solid by following general procedure A with glycine ethyl ester hydrochloride (309 mg), *N*-ethylcyclohexanamine (290 mg) and N-methylpiperazine (652 mg), and the purification was accomplished by chromatography with 10%-20% MeOH in DCM. ^1^**H NMR** (500 MHz, CDCl_3_) δ 5.19 (t, J = 6.0 Hz, 1H), 4.44 – 4.40 (m, 1H), 4.18 (q, J = 7.2 Hz, 2H), 4.07 (d, J = 5.8 Hz, 2H), 3.76 (dd, J = 6.0, 4.4 Hz, 4H), 3.39 (q, *J* = 6.9 Hz, 2H), 2.39 (*t, J* = 5.2 Hz, 4H), 2.30 (s, 3H), 1.80 – 1.63 (m, 5H), 1.47 – 1.28 (m, 4H), 1.25 (t, *J* = 7.2 Hz, 3H), 1.15 – 1.05 (m, 4H).^13^ **C NMR** (126 MHz, CDCl_3_) δ 171.04, 165.84, 164.94, 164.48, 60.87, 54.91, 54.29, 46.14, 43.05, 42.80, 37.04, 30.94, 26.14, 25.77, 15.16, 14.17. LC-MS (ES+): m/z 406.79 [M+H]^+^

Compound **2d** (ethyl (4-(cyclohexyl(ethyl)amino)-6-(4-(2-methoxyphenyl)piperazin-1-yl)-1,3,5-triazin-2-yl)glycinate). **2d** was obtained (538 mg, 1.08 mmol, 50%) as white solid by following general procedure A with glycine ethyl ester hydrochloride (309 mg), N-ethylcyclohexanamine (290 mg) and 1-(2-methoxyphenyl)piperazine (1.25 g), and the purification was accomplished by chromatography with 10%-50% ethyl acetate in hexanes. ^1^**H NMR** (500 MHz, CDCl_3_) δ 7.01 (tt, J = 6.3, 2.9 Hz, 1H), 6.93 (q, *J* = 4.7 Hz, 2H), 6.88 (d, *J* = 7.9 Hz, 1H), 5.20 (t, *J* = 5.9 Hz, 1H), 4.47 (t, *J* = 11.7 Hz, 1H), 4.20 (q, *J* = 7.2 Hz, 2H), 4.10 (d, *J* = 5.8 Hz, 2H), 3.96 – 3.91 (m, 4H), 3.88 (s, 3H), 3.43 (q, *J* = 6.9 Hz, 2H), 3.08 – 3.02 (m, 4H), 1.84 – 1.63 (m, 5H), 1.50 – 1.30 (m, 4H), 1.27 (t, *J* = 7.2 Hz, 3H), 1.19 – 1.08 (m, 4H). ^13^**C NMR** (126 MHz, CDCl_3_) δ 171.17, 166.00, 165.08, 164.63, 152.33, 141.46, 123.10, 121.02, 118.37, 111.22, 60.97, 55.40, 54.38, 50.83, 43.29, 43.17, 37.11, 31.04, 26.23, 25.86, 15.28, 14.27. LC-MS (ES+): m/z 499.00 [M+H]^+^

Compound **3d** (sodium (4-(cyclohexyl(ethyl)amino)-6-(4-(2-methoxyphenyl)piperazin-1-yl)-1,3,5-triazin-2-yl)glycinate). **3d** was obtained (83 mg, 0.169 mmol, 84%) as light-yellow powder by following general procedure B with **2d** (100 mg).^1^ **H NMR** (500 MHz, DMSO) δ 6.98 – 6.91 (m, 2H), 6.86 (qd, J = 7.8, 2.4 Hz, 2H), 5.87 (d, J = 15.4 Hz, 1H), 4.53 – 4.34 (m, 1H), 3.78 (s, 4H, br), 3.65 – 3.48 (m, 4H), 2.92 (s, 4H), 1.75 (d, J = 12.4 Hz, 2H), 1.65 – 1.56 (m, 3H), 1.51 – 1.39 (m, 2H), 1.34 – 1.23 (m, 2H), 1.14 – 1.06 (m, 4H). ^13^**C NMR** (126 MHz, DMSO) δ 173.11, 164.93, 164.77, 164.24, 152.08, 141.23, 122.66, 120.82, 118.22, 111.84, 56.03, 55.32, 53.72, 50.21, 42.89, 36.67, 30.51, 25.89, 25.28, 15.41. LC-MS (ES+): m/z 470.90 [M-Na+2H]^+^

## Acknowledgments

We acknowledge funding support by R01CA293456 (PI Gabr) from the National Cancer Institute (NCI).

## Data availability

The authors confirm that all data are available as ESI.† Furthermore, additional data and original files are available from the authors upon request.

