## Supporting Information for "Discovery of a First-in-Class SLIT2 Binder Disrupting the SLIT2/ROBO1 Axis via DNA-Encoded Library (DEL) Screening"

*Electronic Supplementary Information*

| **Contents** |  |
| --- | --- |
| ^1^H NMR spectrum of compound **DEL-S1** | S2 |
| LCMS report of compound **DEL-S1** | S3 |
| Mass spectrum of compound **DEL-S1** | S4 |
| HPLC purity of compound **DEL-S1** | S5 |
| Chiral SFC report of compound **DEL-S1** | S6 |
| Cell viability as assessed by PrestoBlue of NHA **(A)**, hBMECs **(B)**, and HepG2 **(C)** upon incubation with a single dose of 50 μM of **DEL-S1** after 72 h incubation. | S7 |
| Characterization of newly synthesized compounds | S8 |

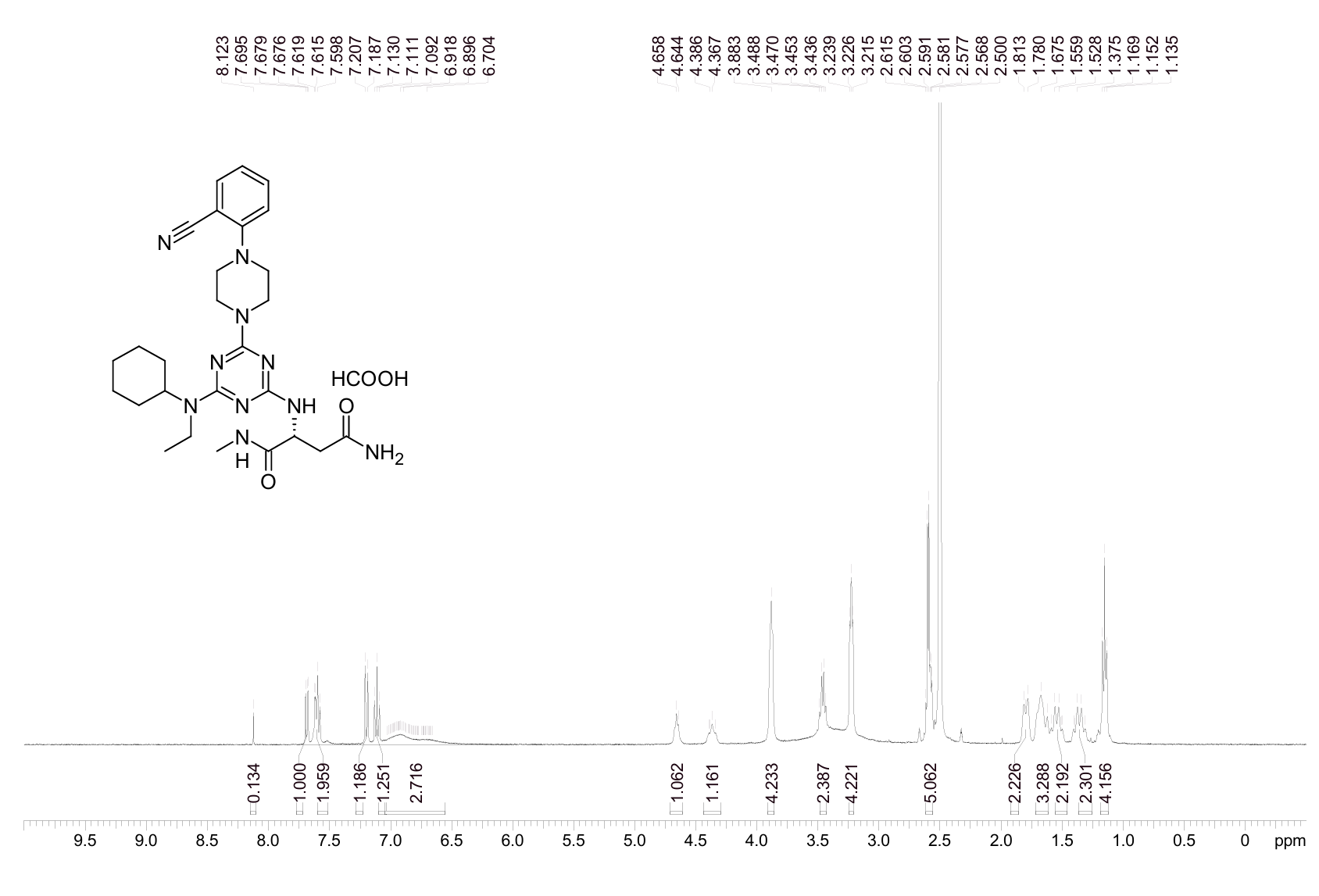

**Figure S1**. ^1^H NMR spectrum of compound **DEL-S1**.

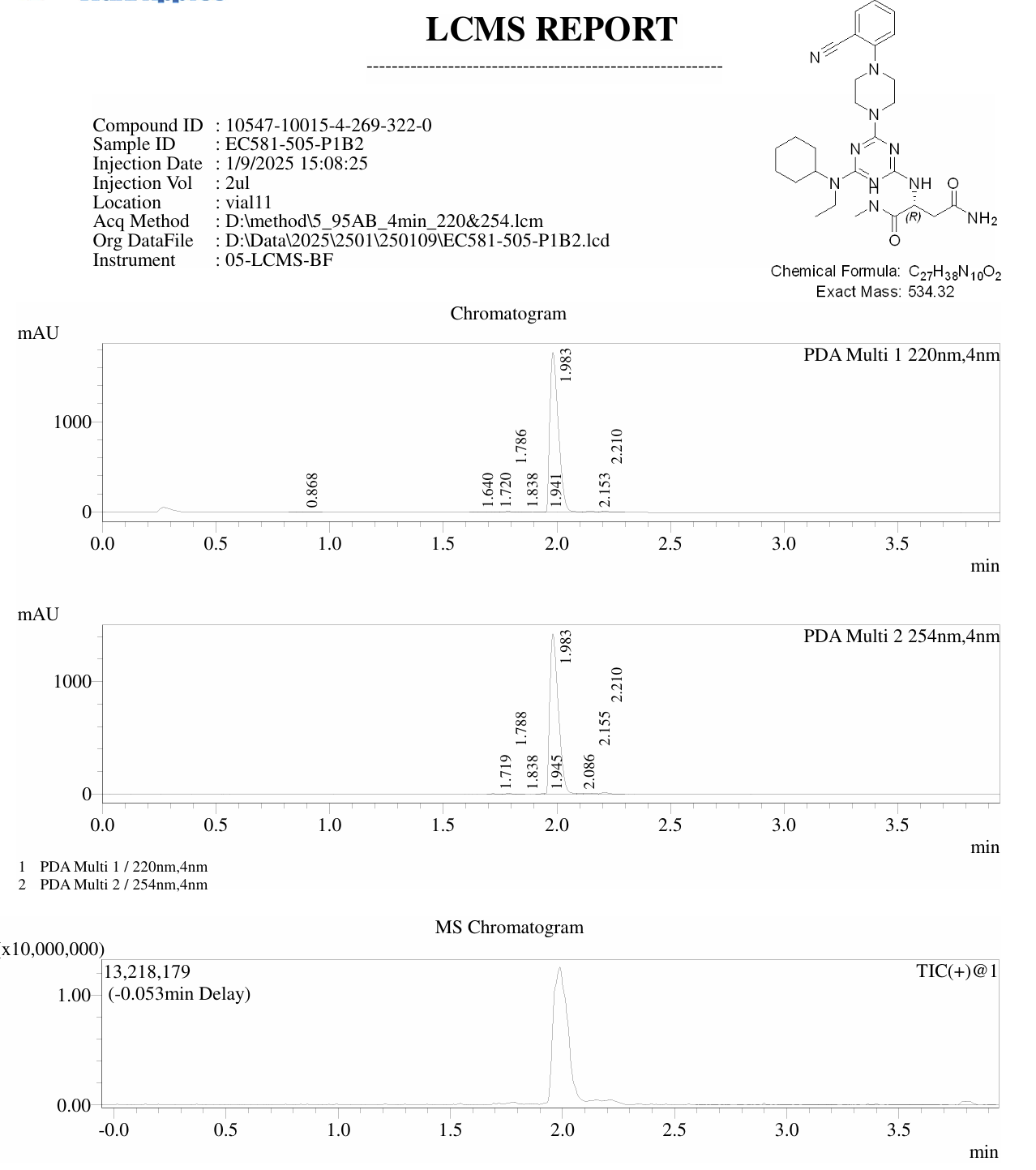

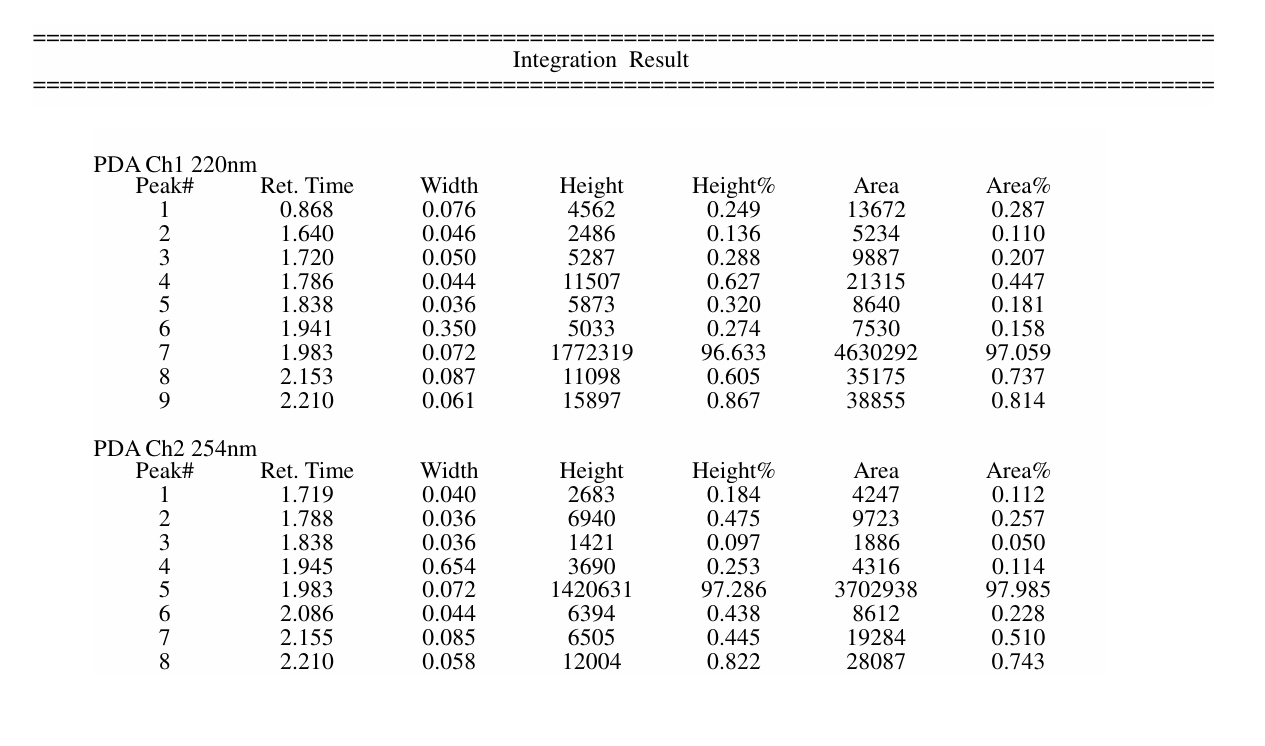

**Figure S2**. LCMS report of compound **DEL-S1**.

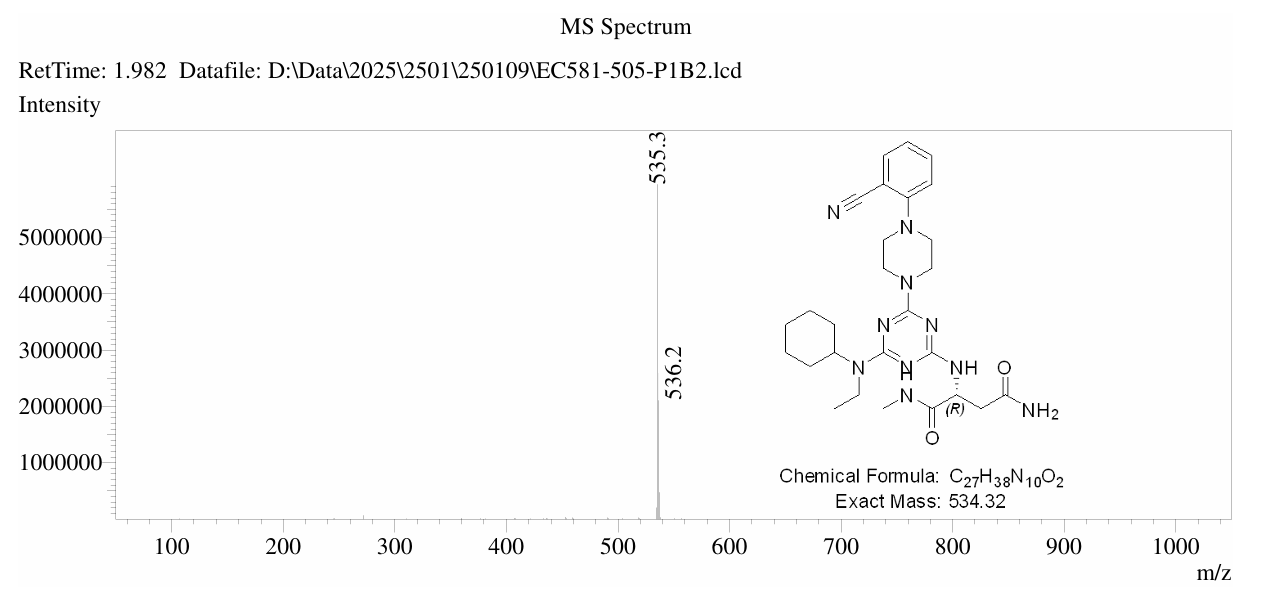

**Figure S3**. Mass spectrum of compound **DEL-S1**.

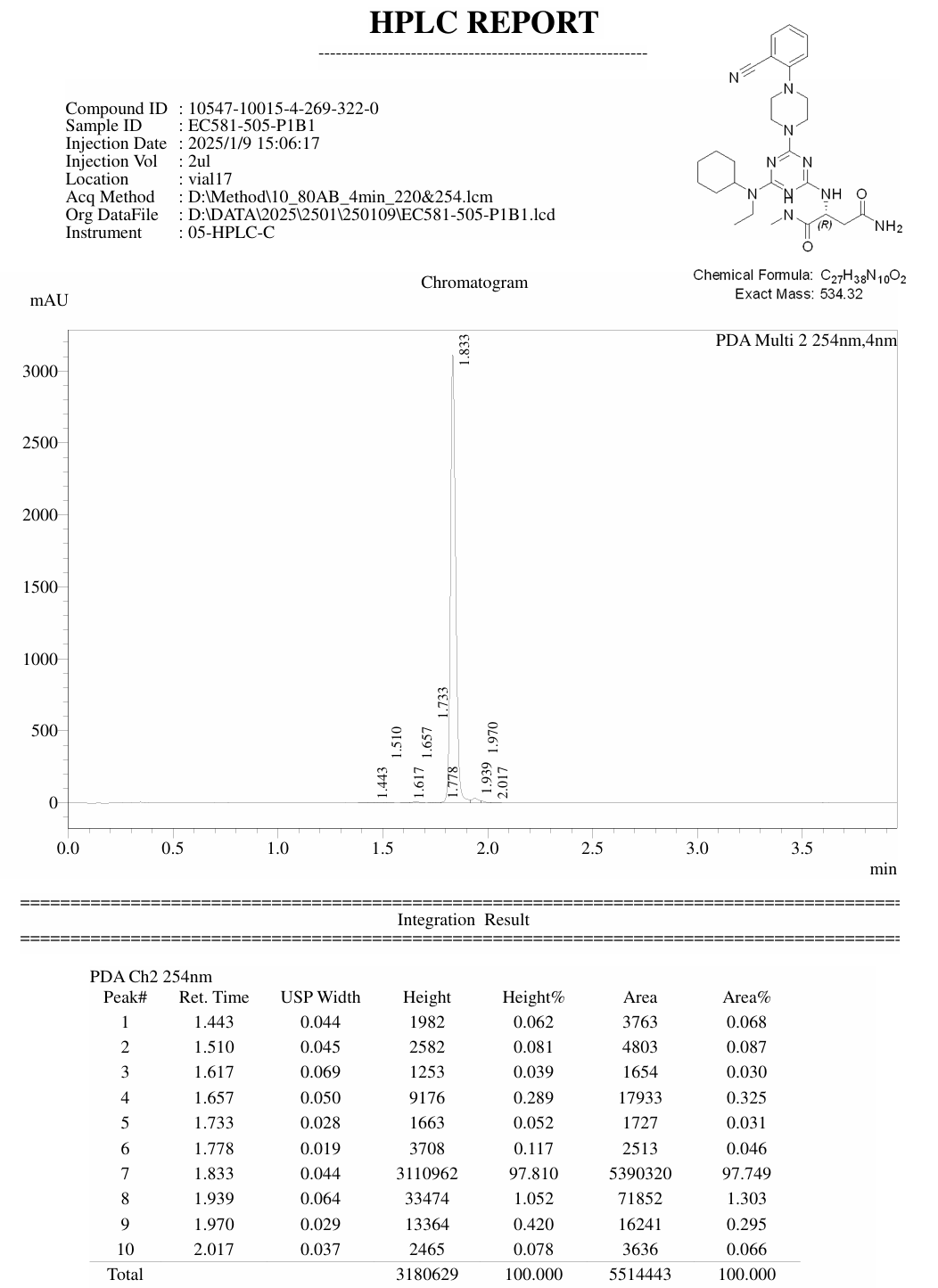

**Figure S4**. HPLC purity of compound **DEL-S1**.

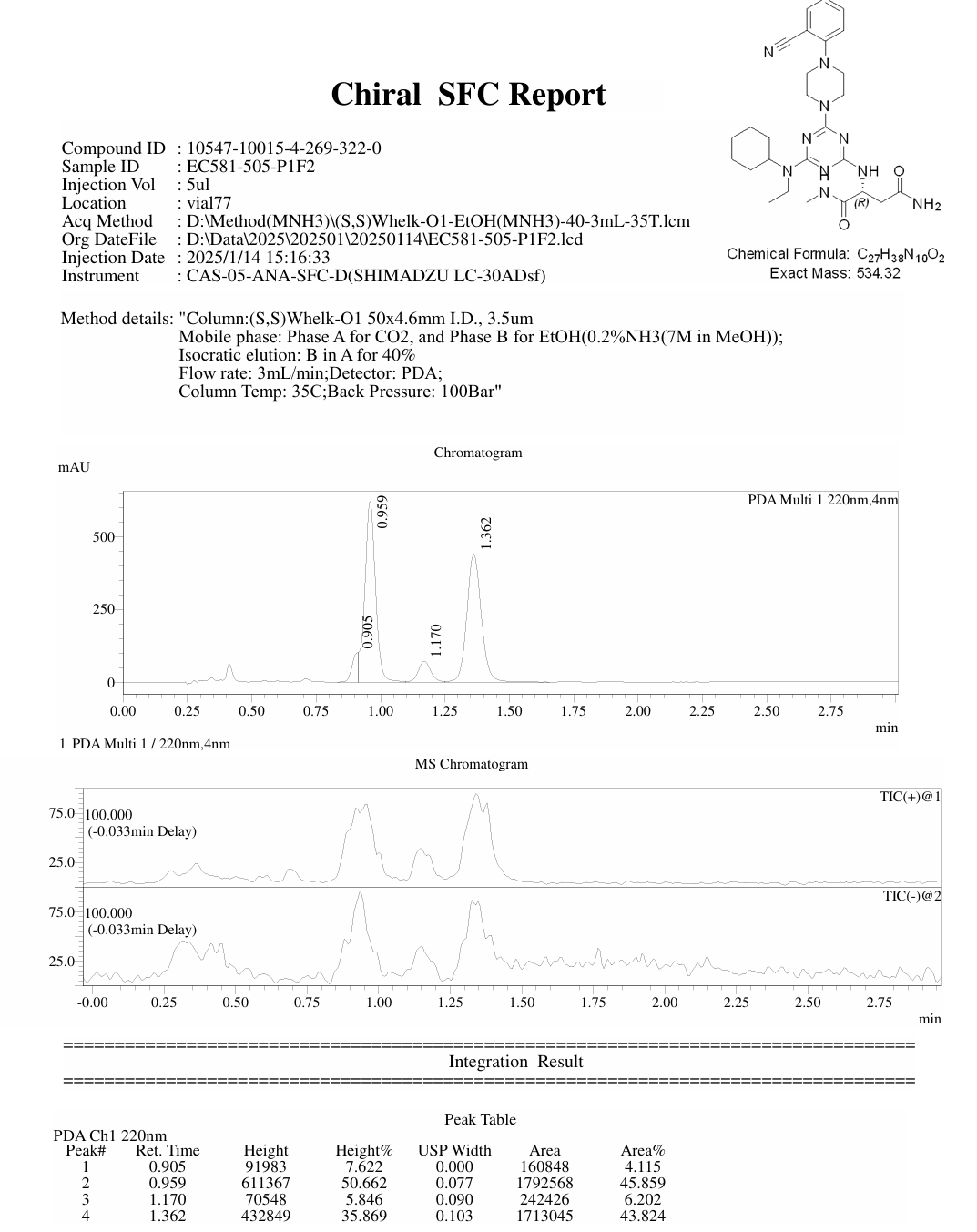

**Figure S5**. Chiral SFC report of compound **DEL-S1**.

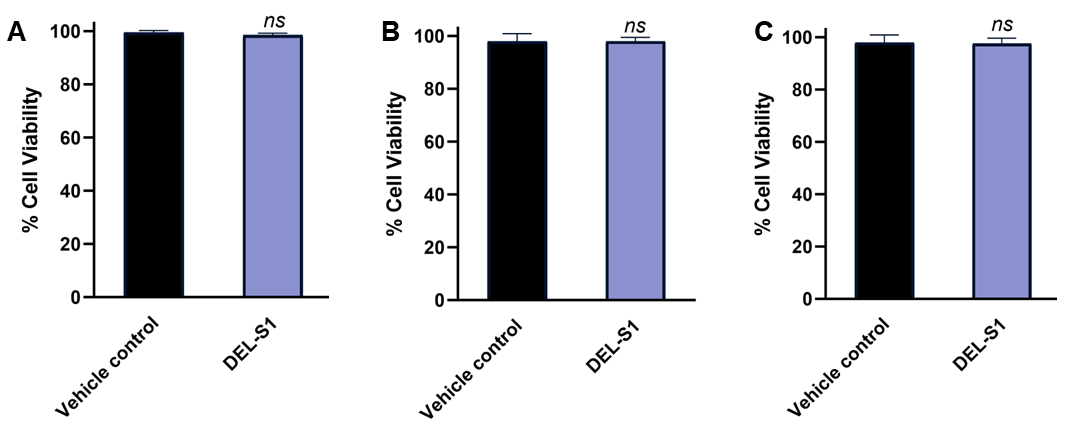

**Figure S6**. Cell viability as assessed by PrestoBlue of NHA **(A)**, hBMECs **(B)**, and HepG2 **(C)** upon incubation with a single dose of 50 μM of **DEL-S1** after 72 h incubation. *ns* denotes nonsignificant relative to vehicle control. Error bars represent standard deviation (n = 3).

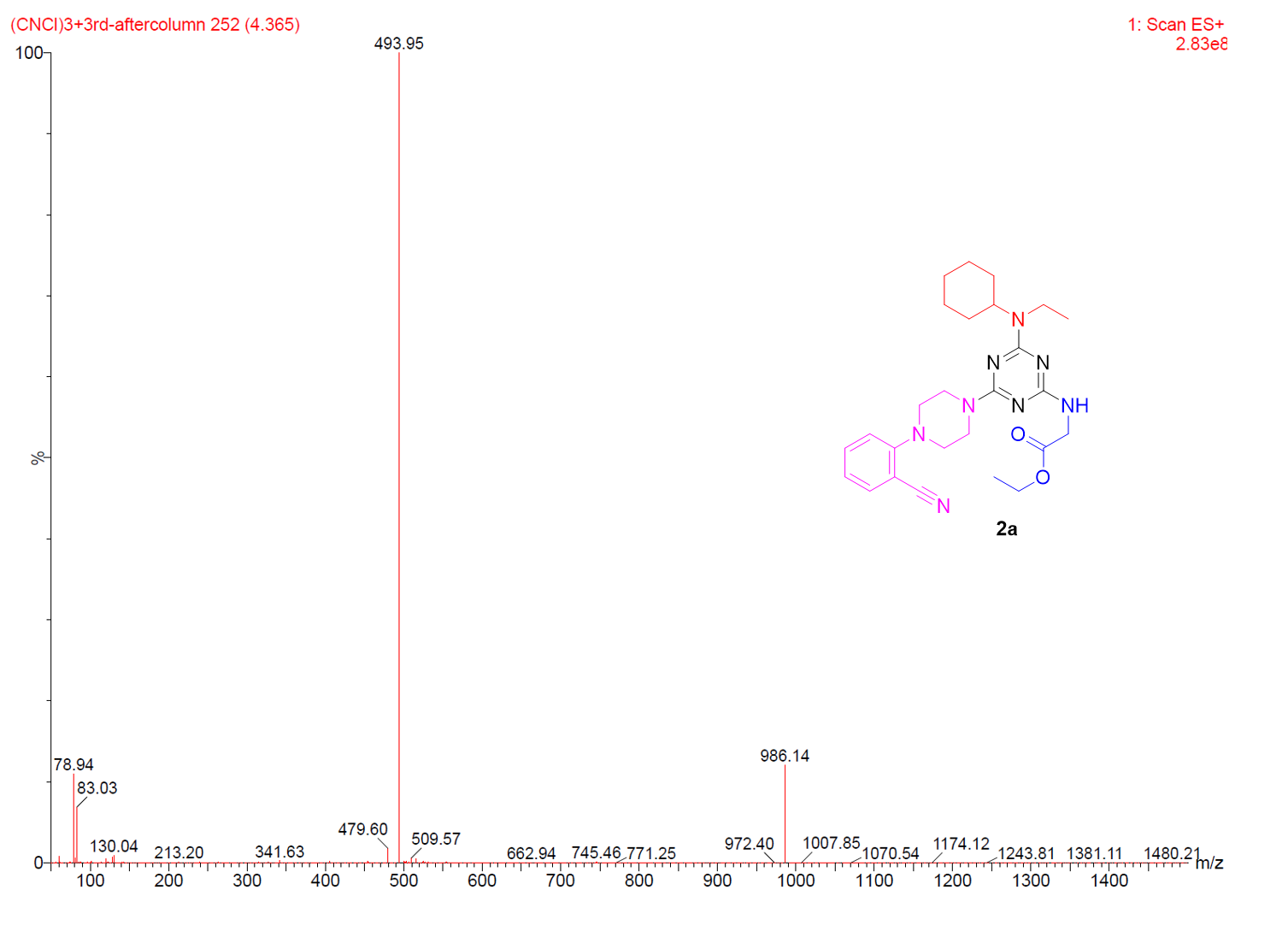

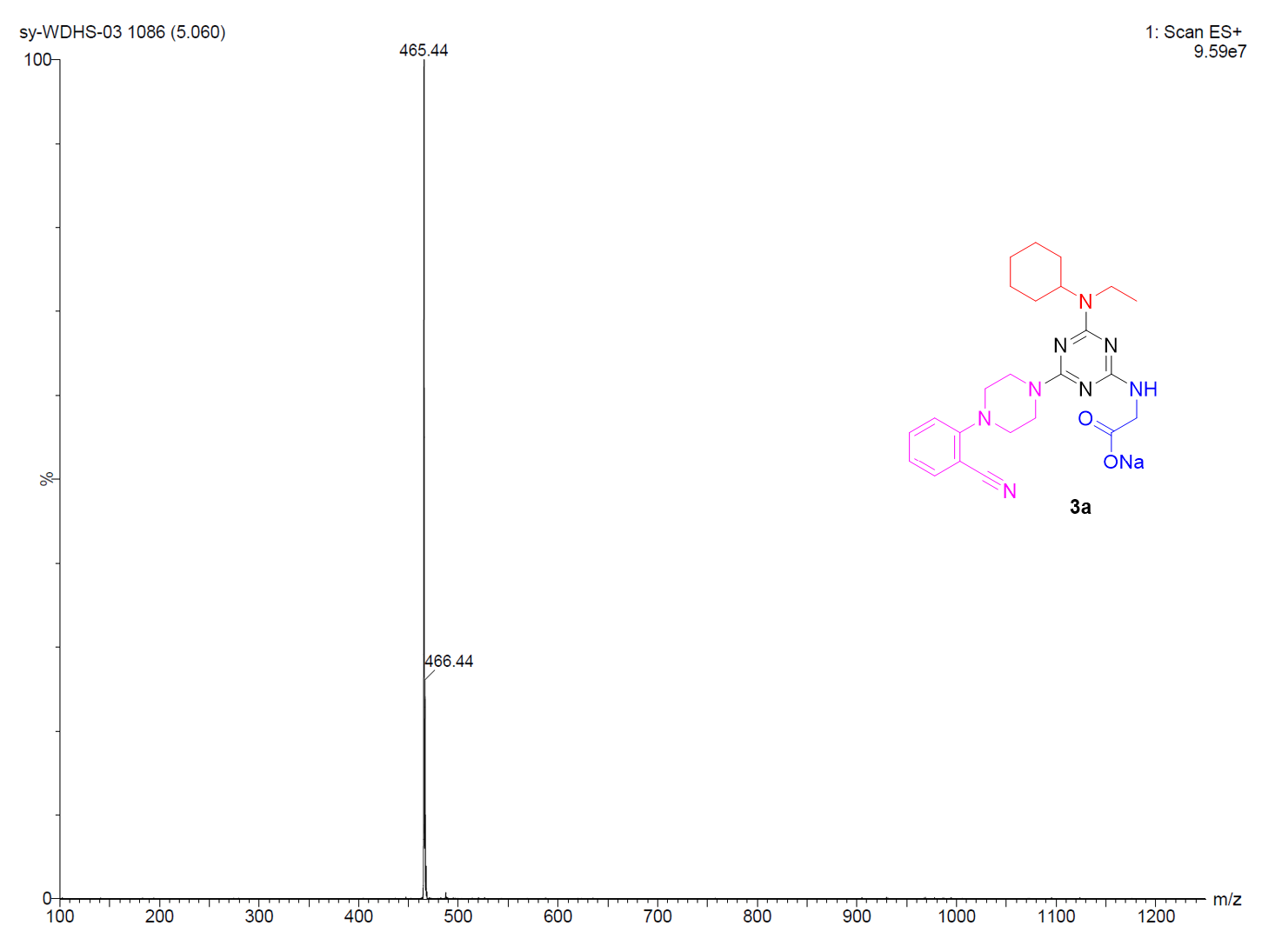

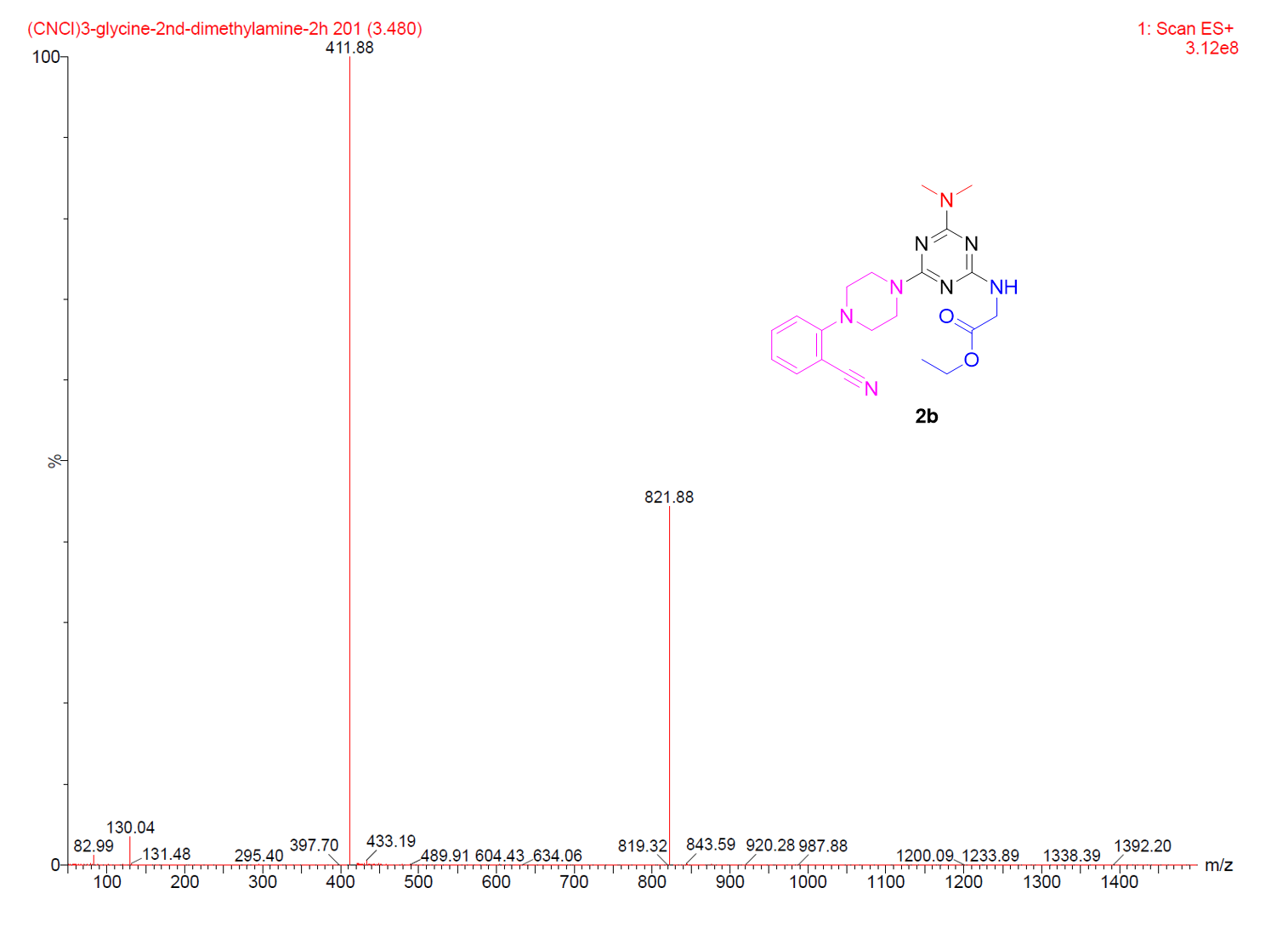

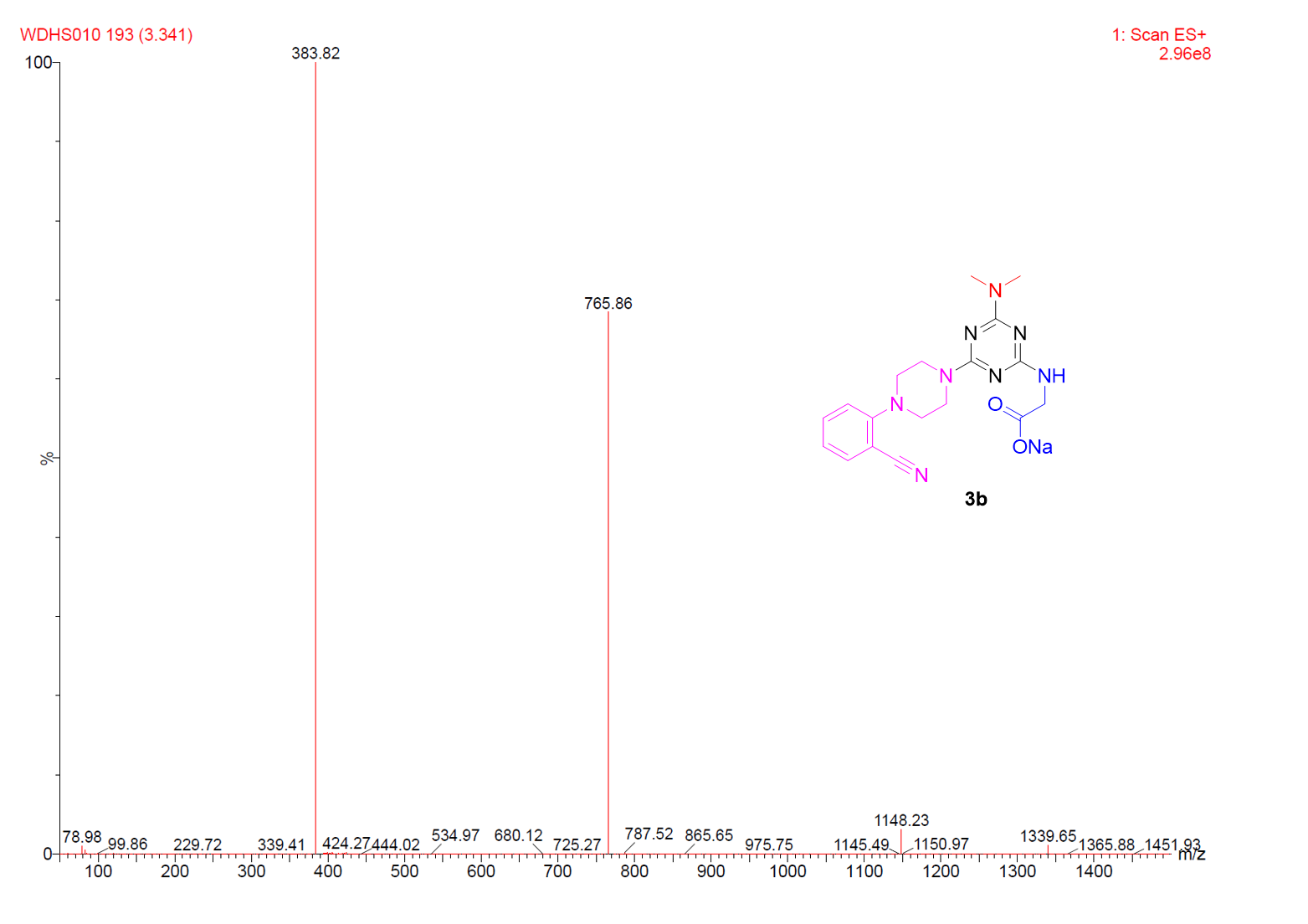

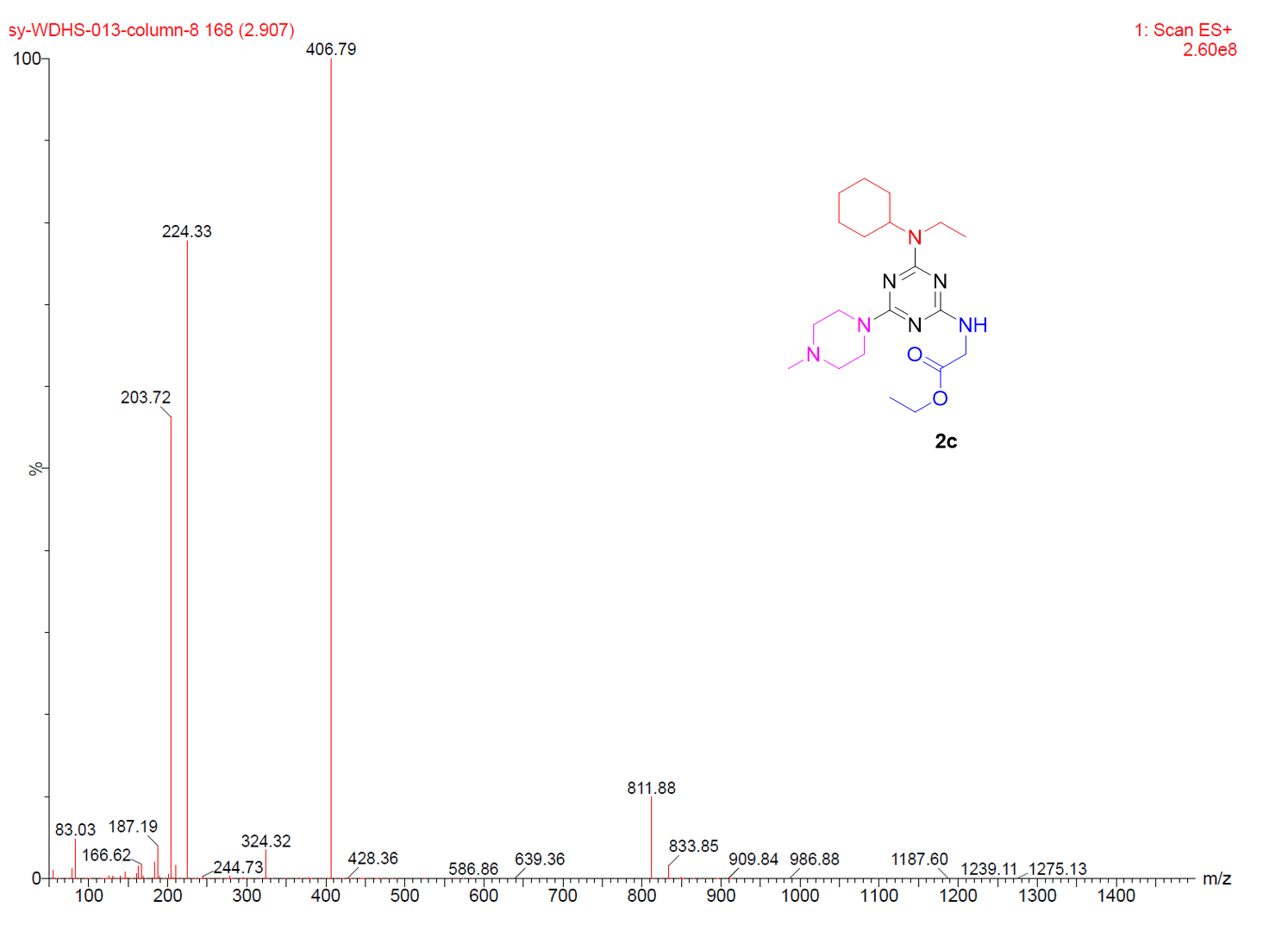

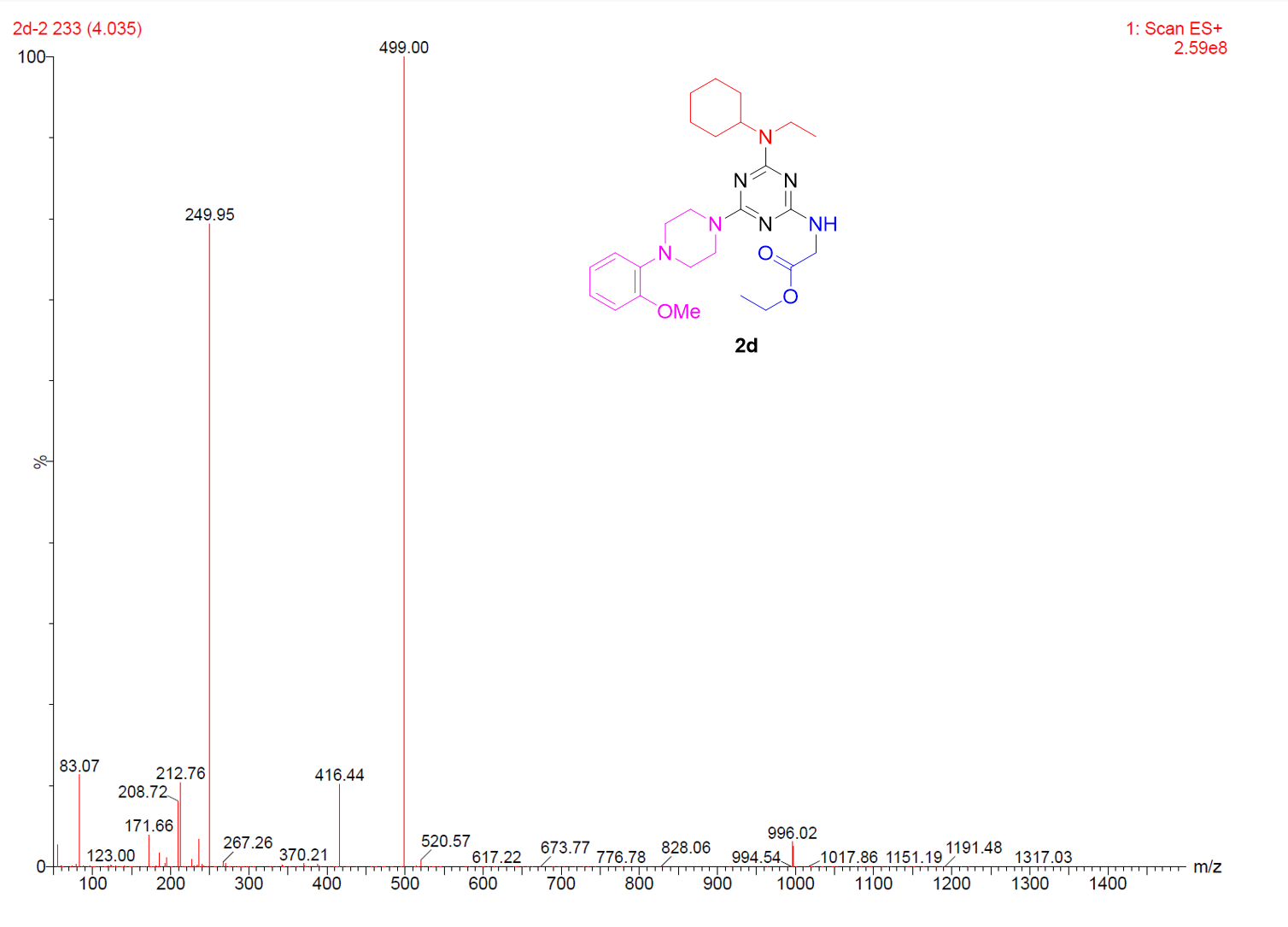

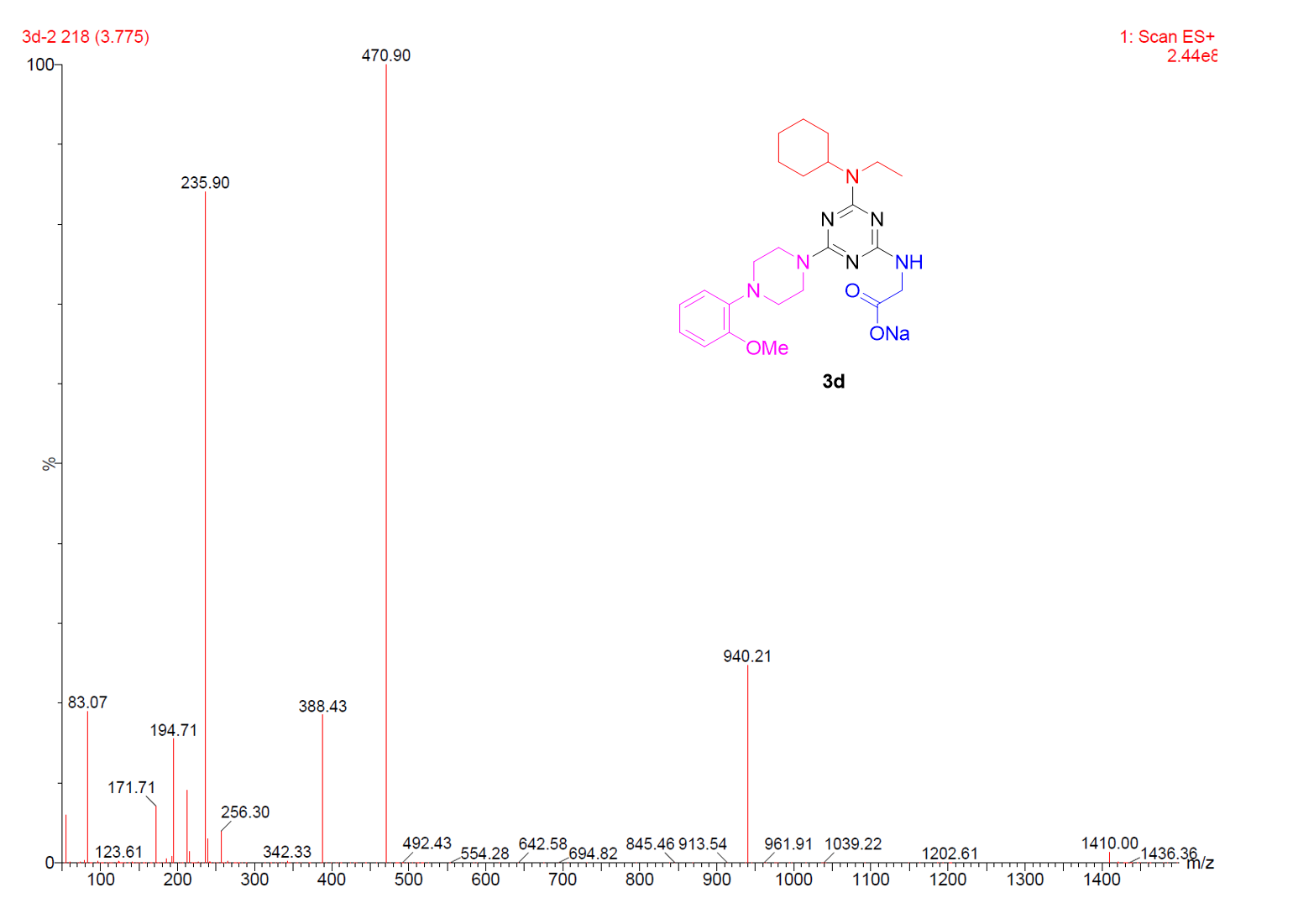

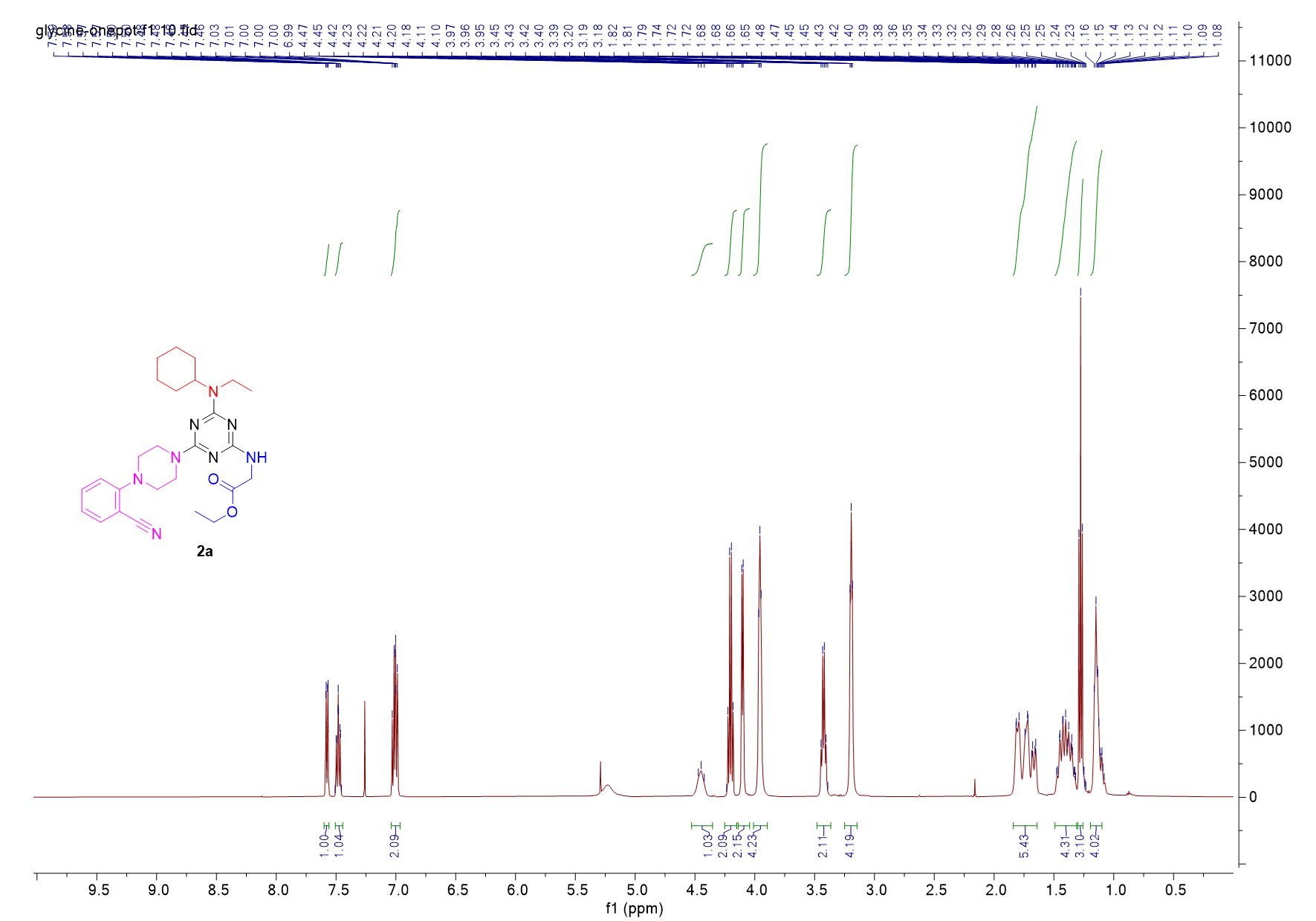

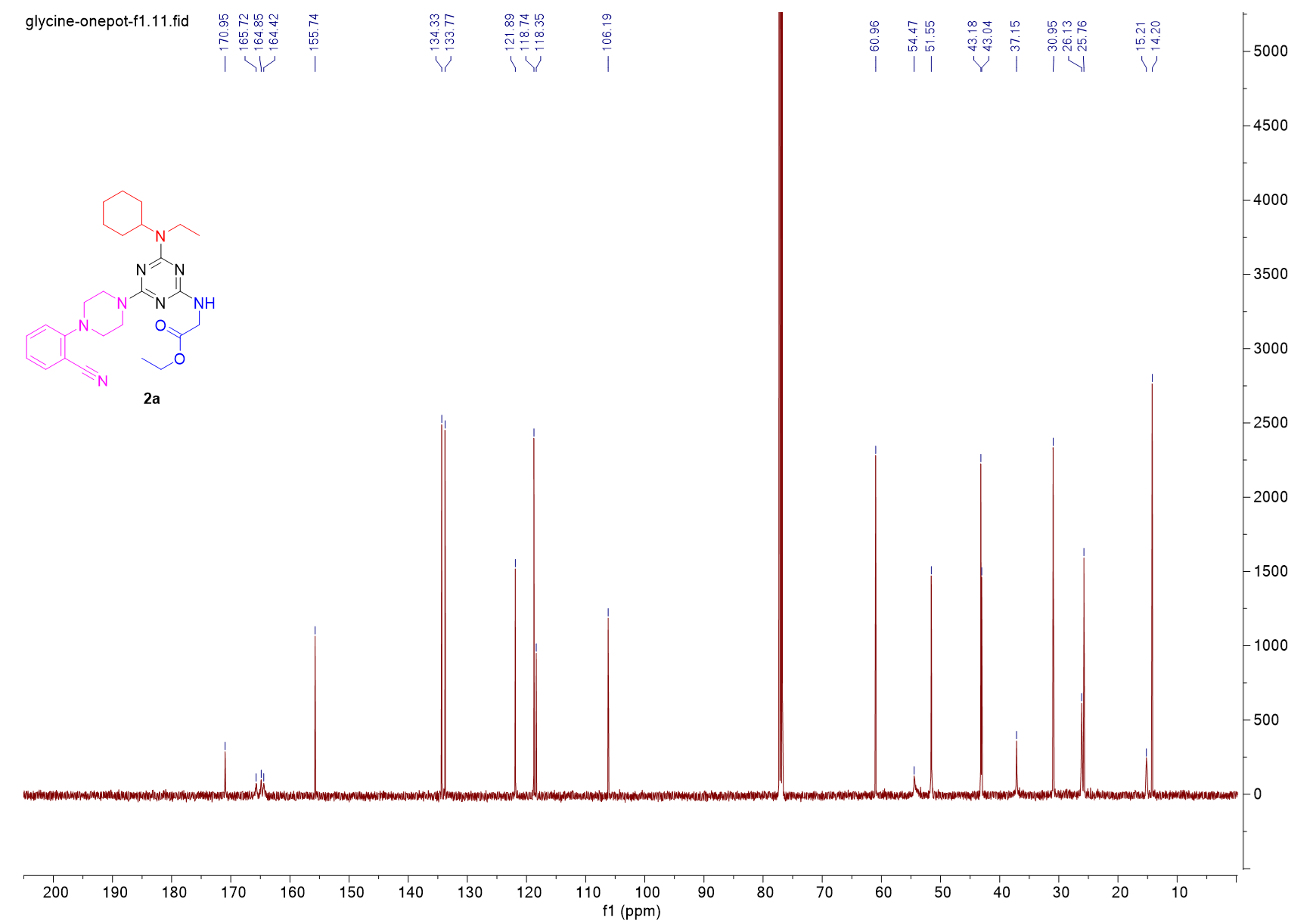

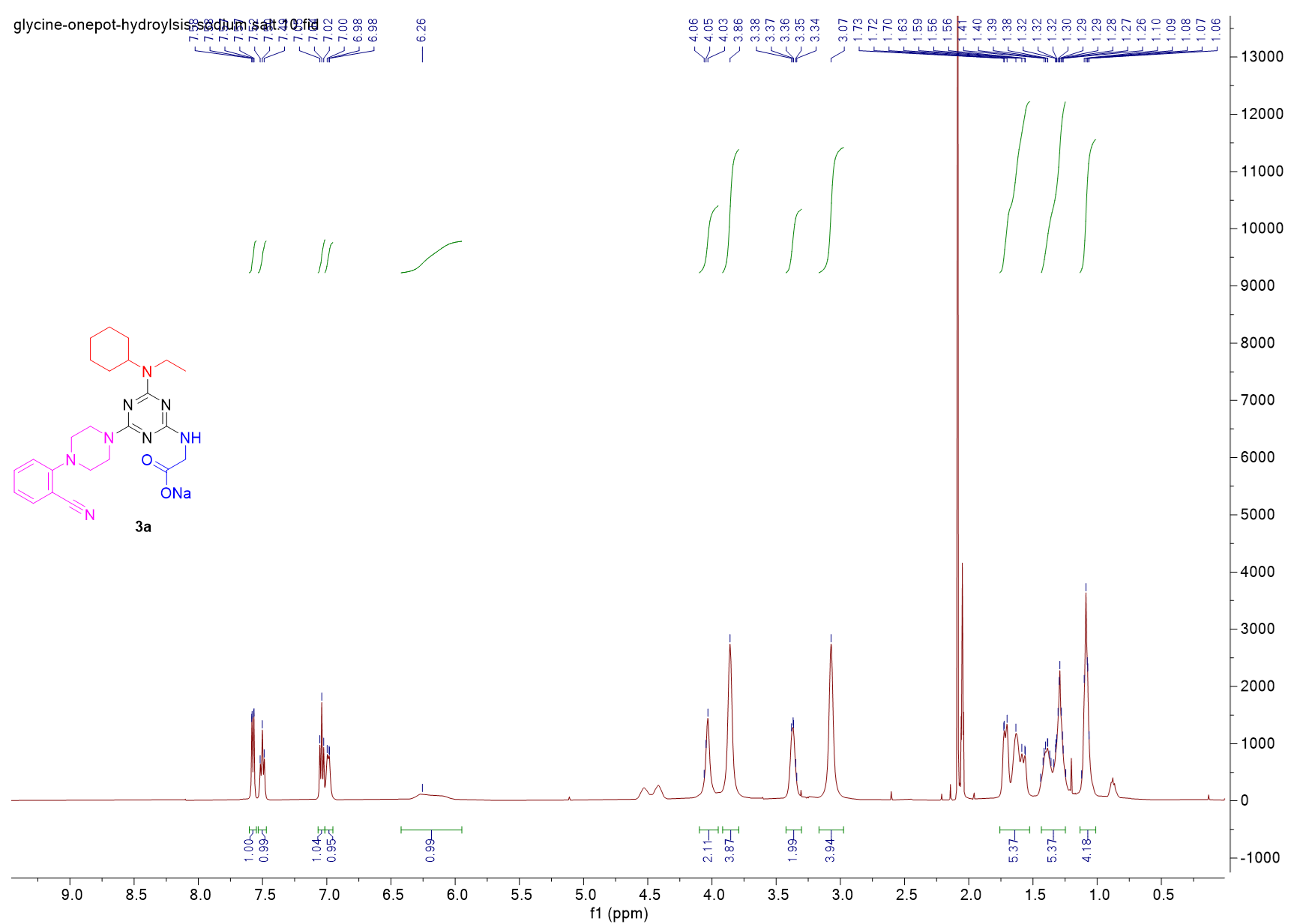

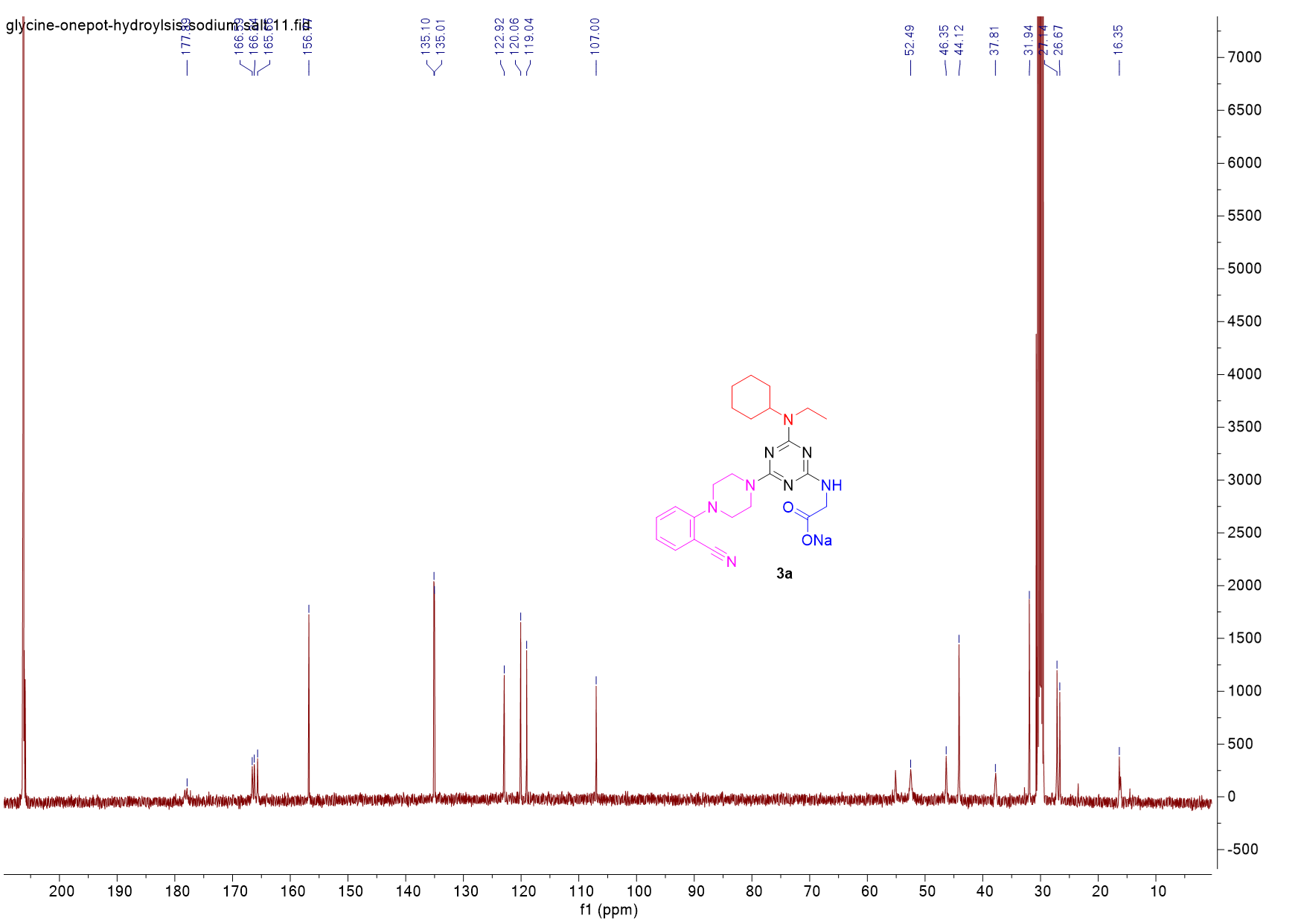

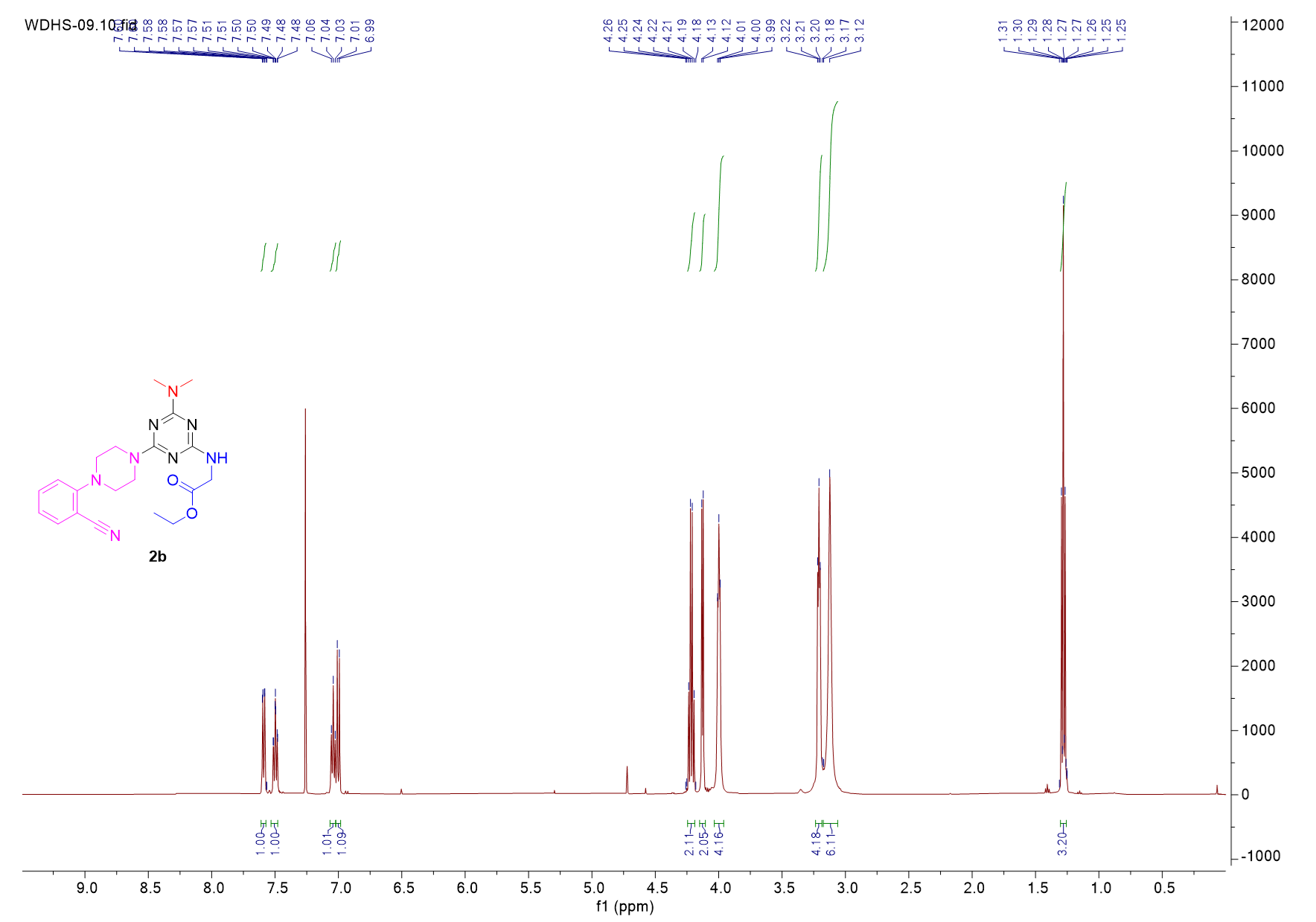

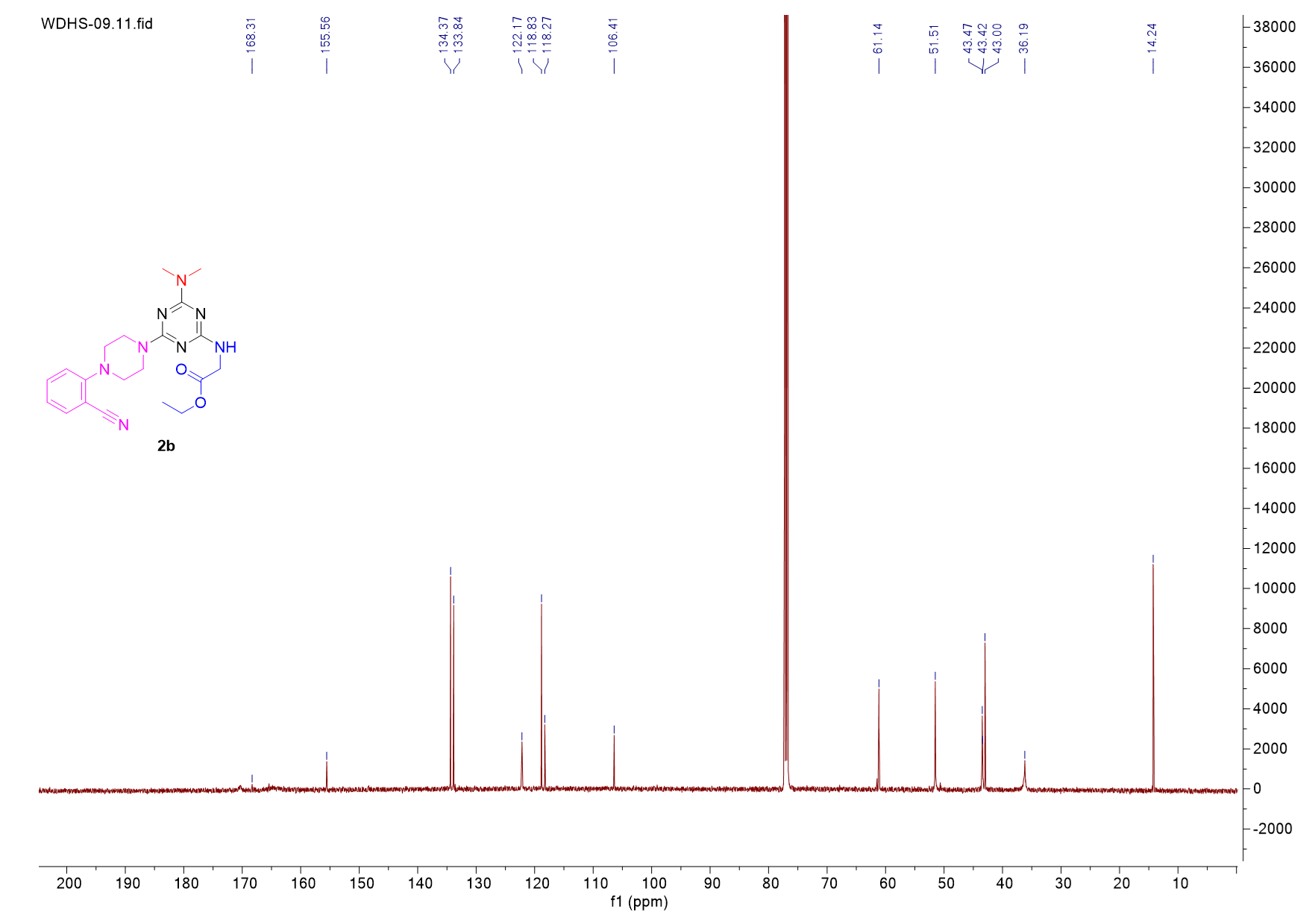

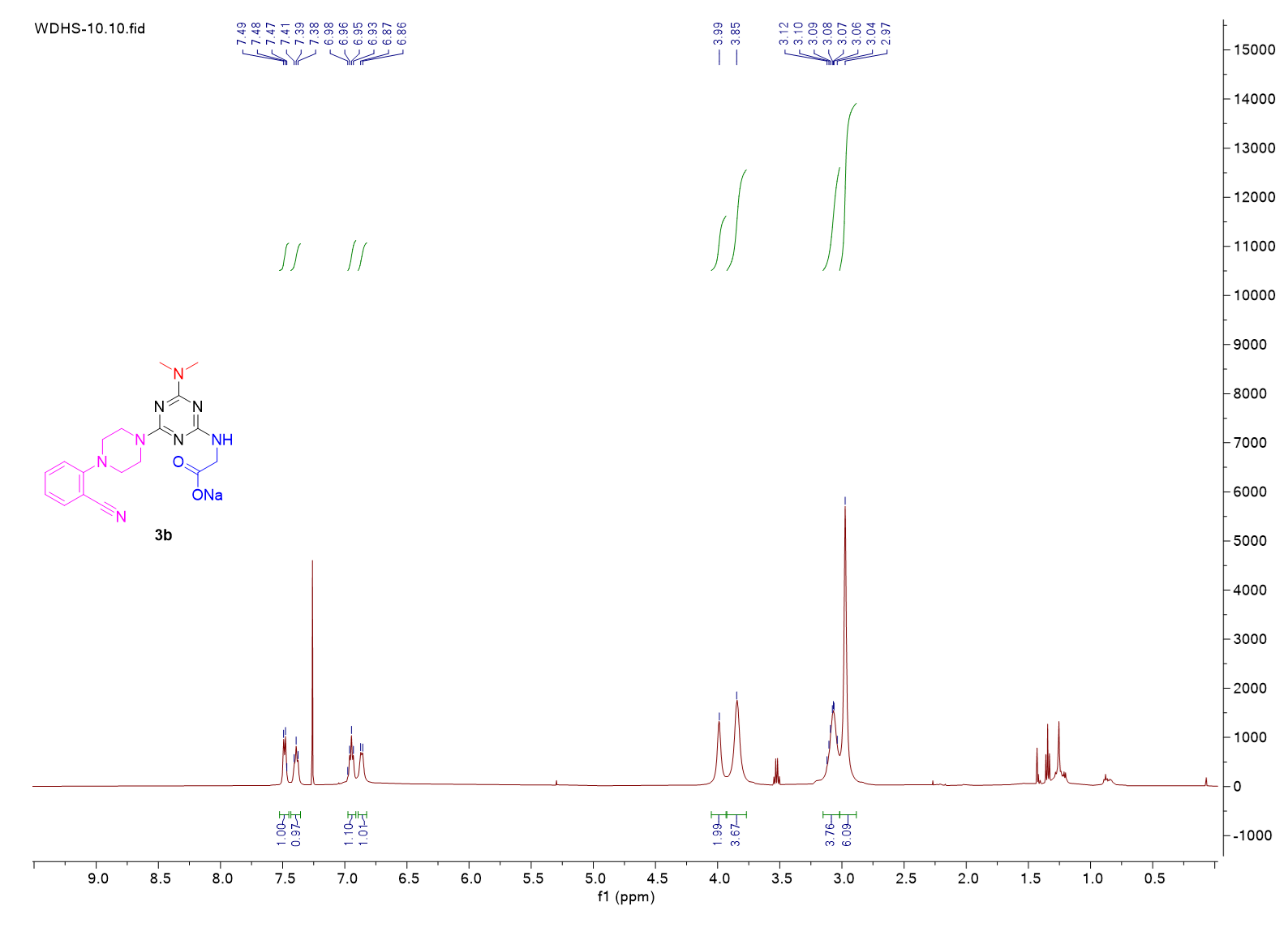

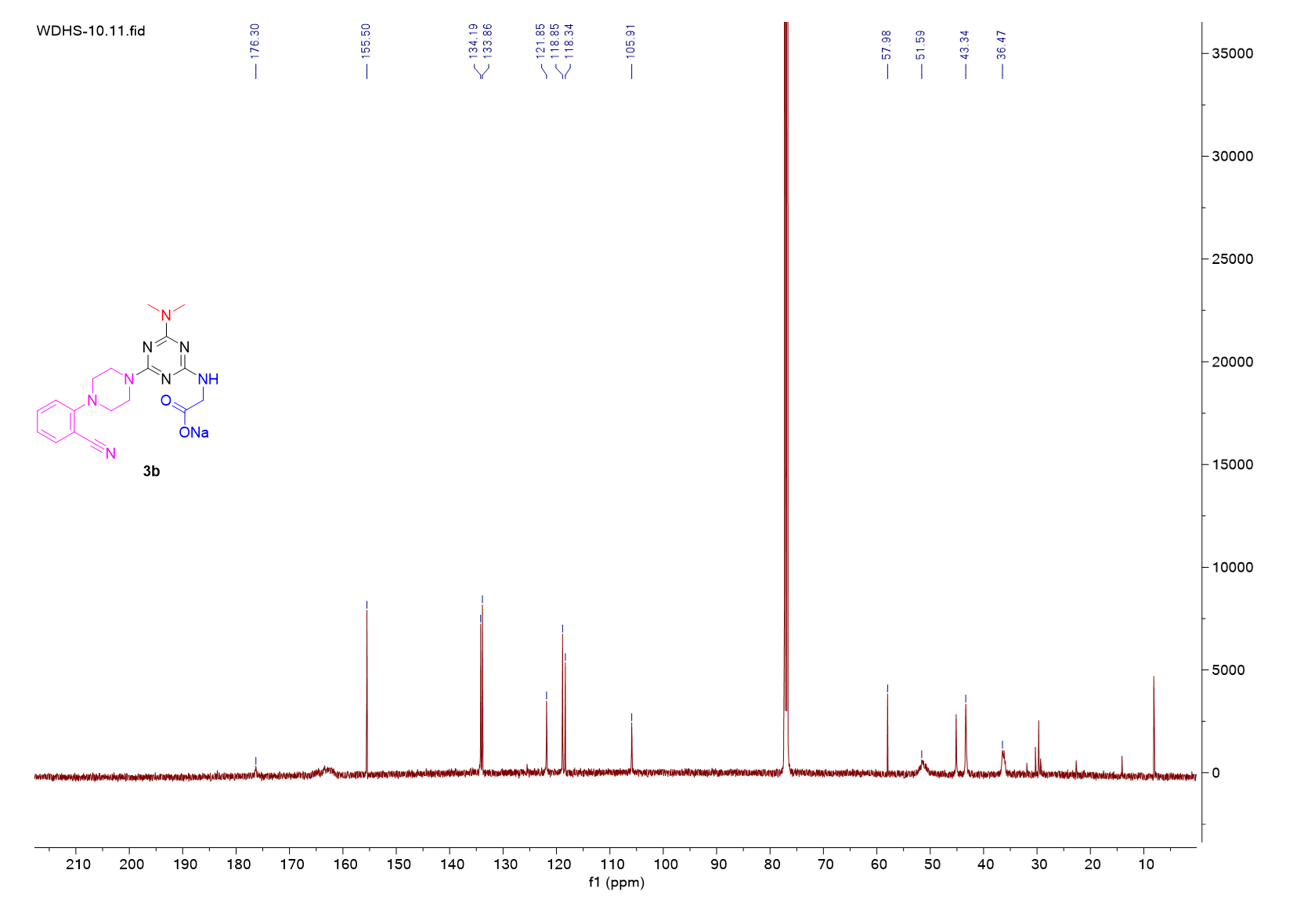

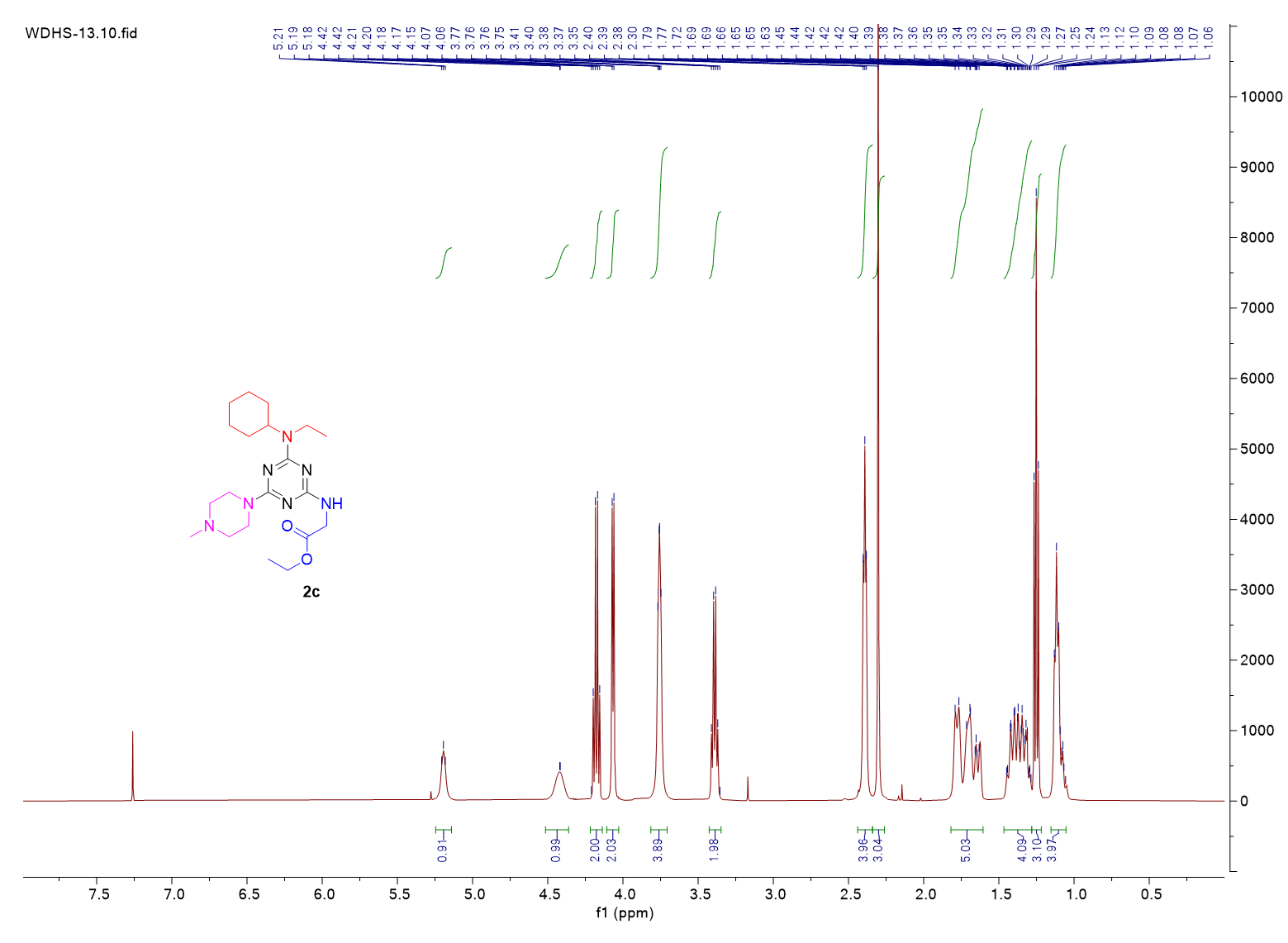

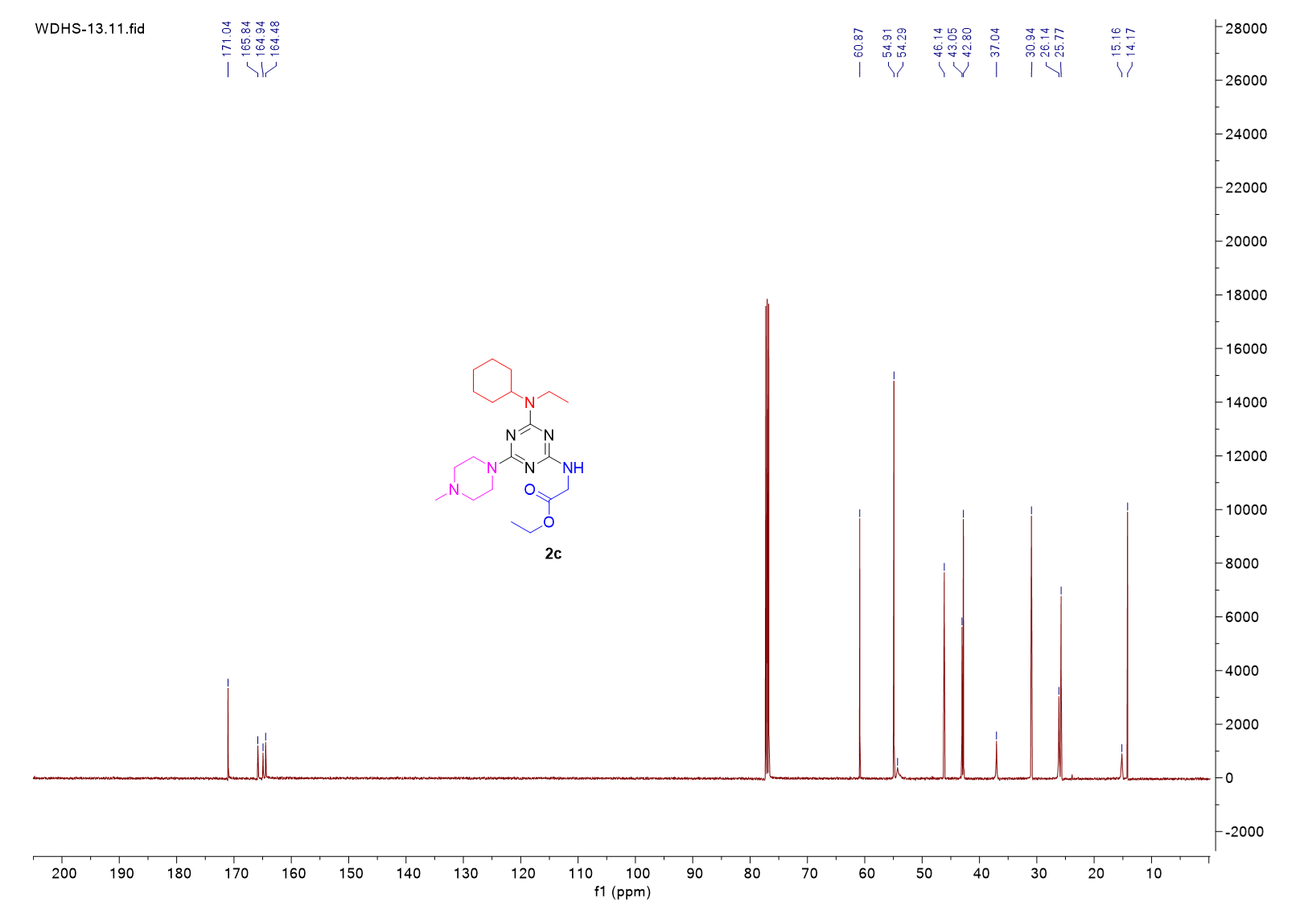

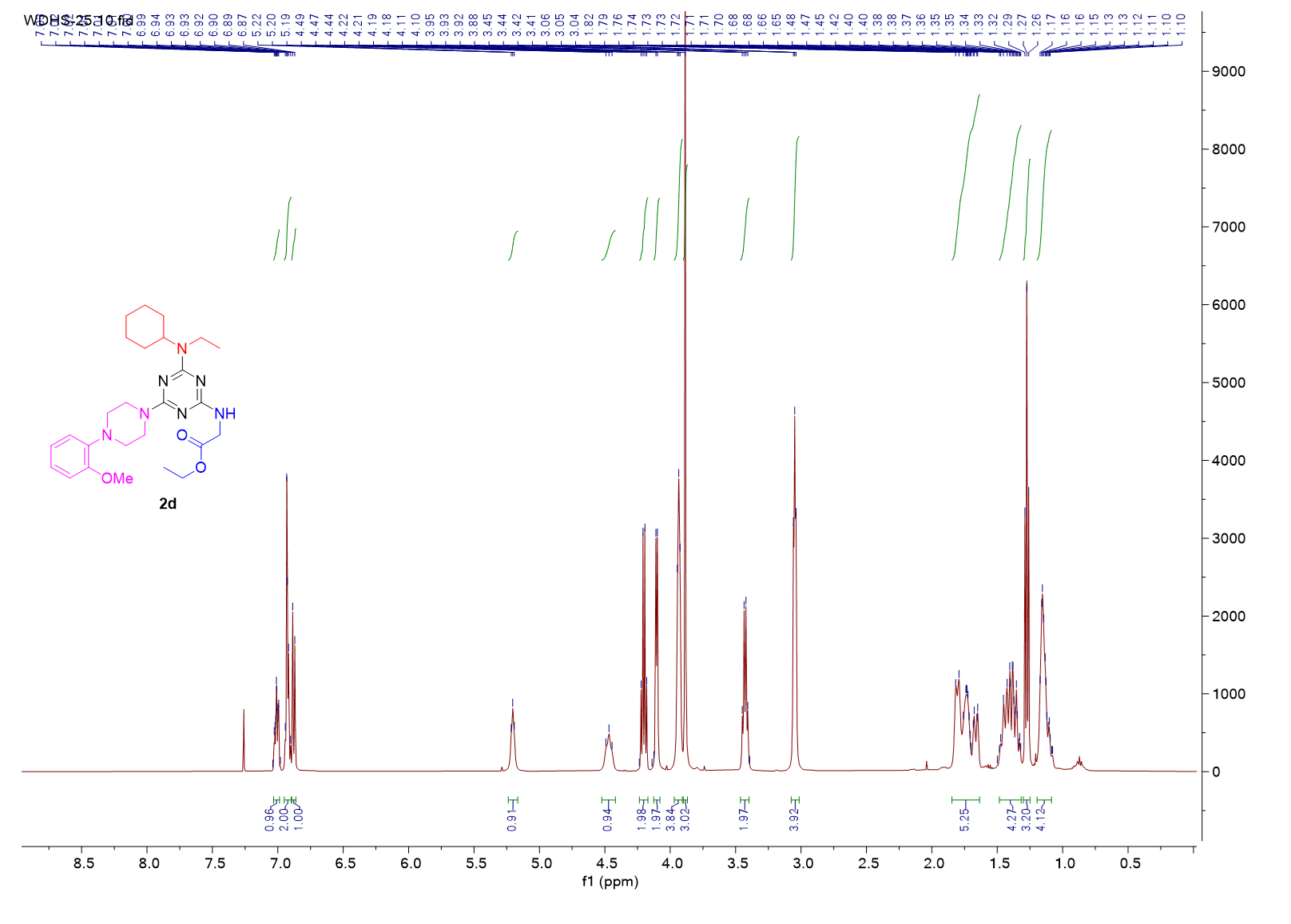

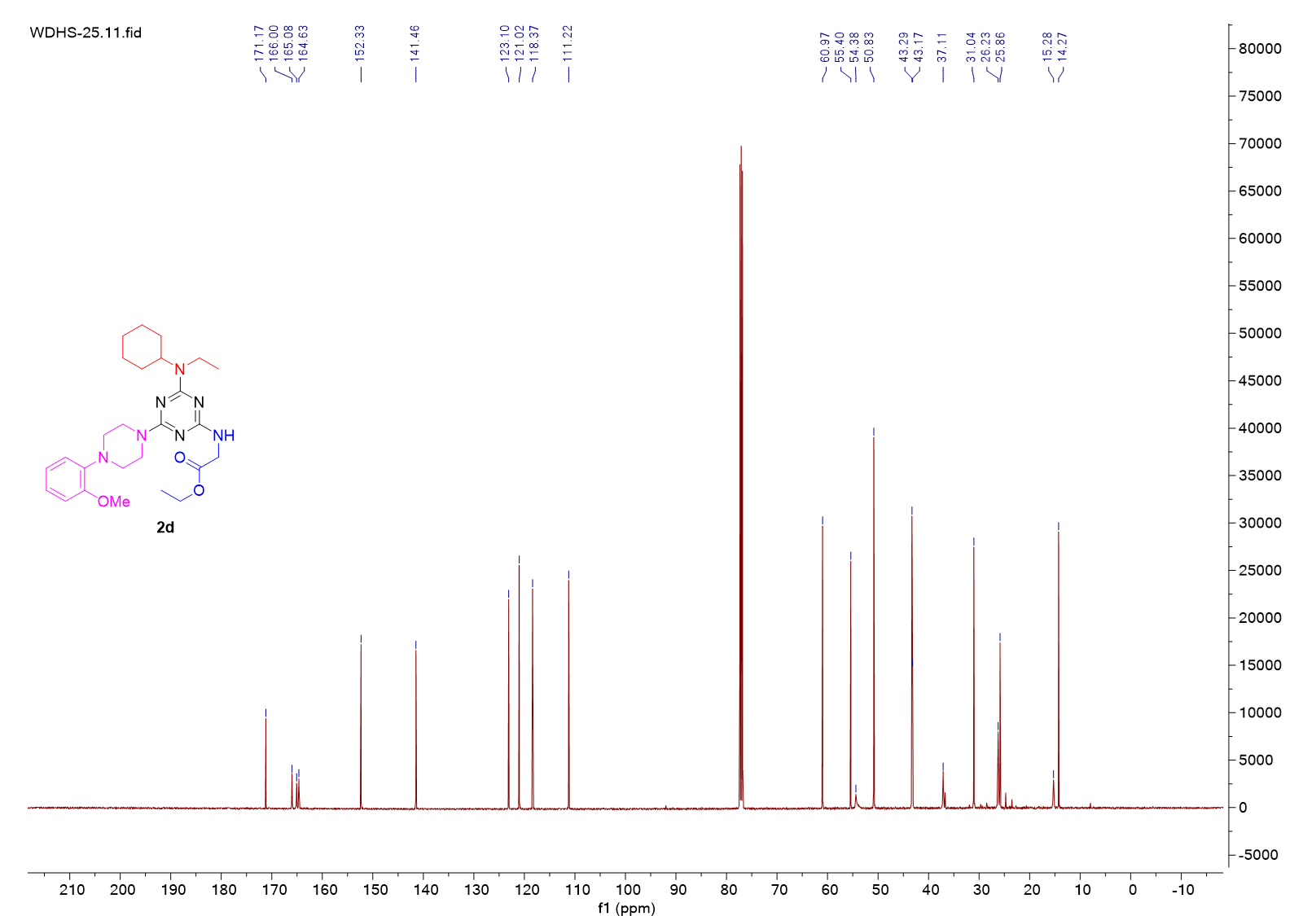

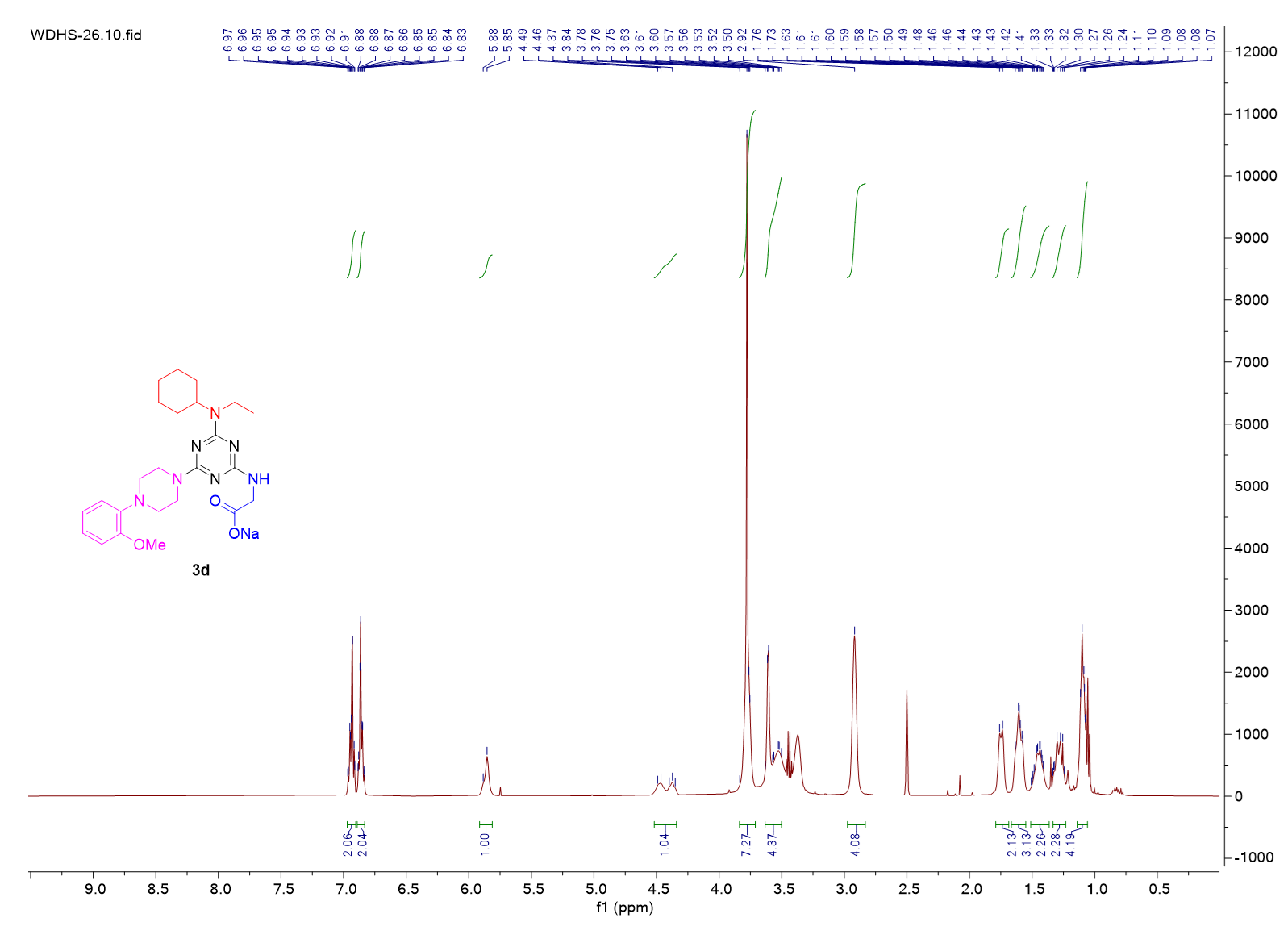

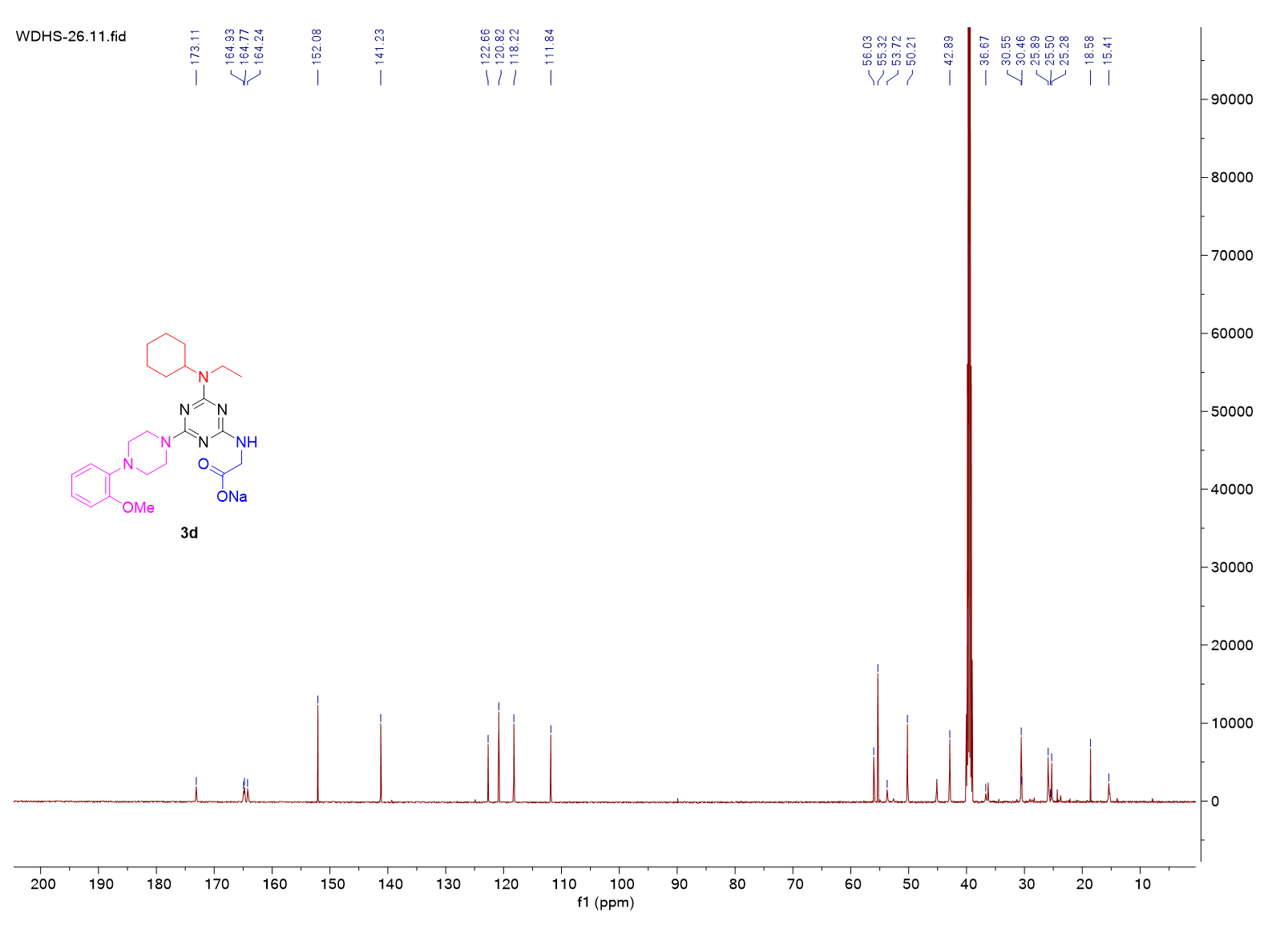
